# Third-nucleotide codon bias and synonymous codon bias define functional translational programs that shape human tissue and cancer proteomes

**DOI:** 10.1101/2025.10.31.685942

**Authors:** Sherif Rashad, Kuniyasu Niizuma

## Abstract

**Background:** Codon usage bias is a universal feature of the genetic code, yet how synonymous codon bias or third-nucleotide codon bias (A/T-vs G/C-ending) shape translation and proteome composition across tissues and cancer remain unclear.

**Results:** Using comparative genomics between human and rodent coding sequences, we uncovered a conserved codon-bias axis. A/T-ending codons consistently marked genes involved in proliferation and RNA processing, whereas G/C-ending codons were enriched for differentiation and neuronal functions. While GC3 scores, measuring the third-nucleotide codon bias, showed differences between humans and rodents due to recombination events, the functional dichotomy was conserved. Isoacceptors frequencies, measuring gene synonymous codon bias, was conserved from rodents to humans. Synonymous codons exhibited distinct functional enrichment patterns, demonstrating functional divergence at the codon level. Two new indices; the ANN-index and m⁷G-index, reflecting codons decoded by the t⁶A and m⁷G tRNA modifications, linked tRNA modification biology to translation. Both indices correlated with proliferative, A/T-biased programs, providing a universal basis for their roles in cancer. Tissue proteomes showed strong RNA–protein discordance and distinct codon biases. Analysis of 21 cancer types revealed a global A/T-ending codon bias in cancer. Analysis of 2,600 cancer cell lines revealed codon bias heterogeneity in cell lines from the same cancer subtype that is not observable between cancer patients.

**Conclusions:** Our results define synonymous codon divergence and tRNA-modification indices as determinants of translational reprogramming. This work establishes a unified framework connecting codon usage, tRNA modifications, and proteome remodeling, providing a basis for rational design of mRNA and gene therapeutics.

**Graphical abstract:** 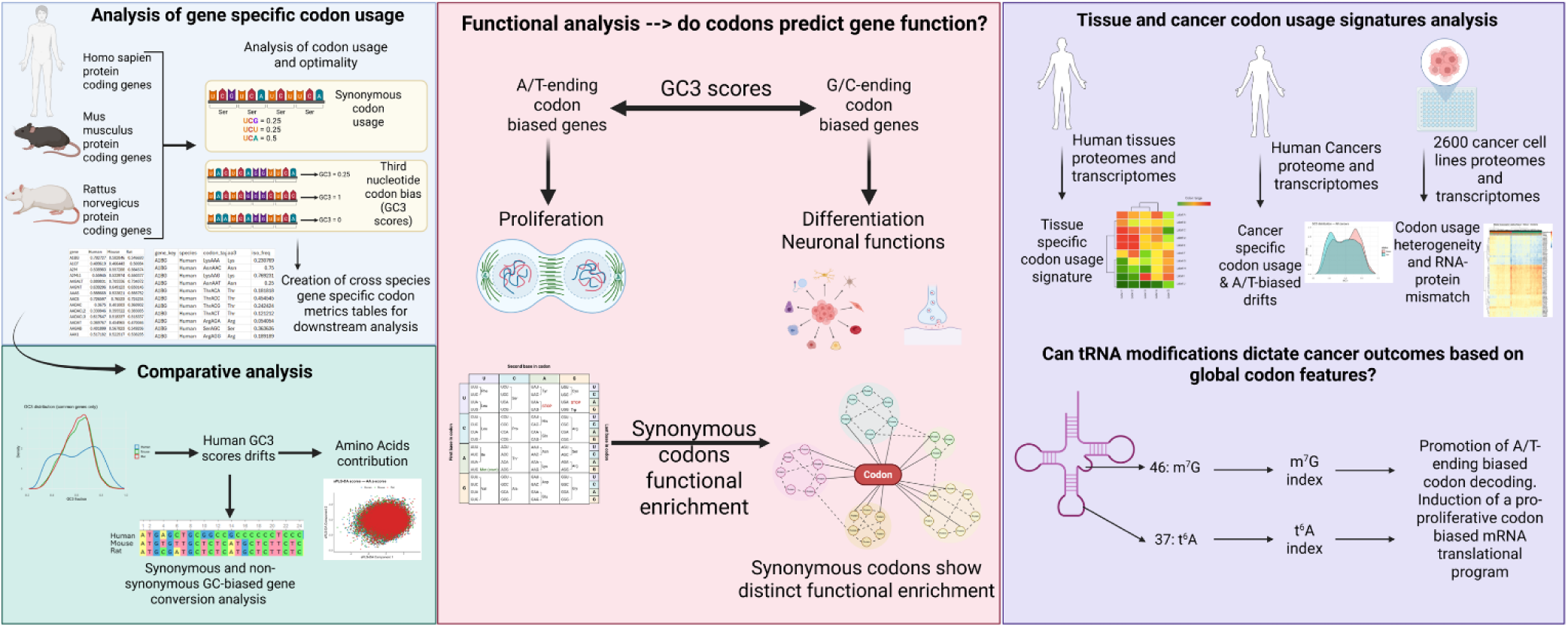

## Background

The translation of the genetic code to protein is a heavily regulated and energy expensive process that entails the translation of genetic information, in the form of codons, stored in the coding sequences of mRNAs to amino acids that form proteins. During mRNA translation, a limited set of tRNA anticodons (48 in human and 47 in mouse) are available to decode the 64 mRNA codons encoding for the 20 essential amino acids (1). This arrangement leads to a degree of genetic code redundancy, where most amino acids (apart from tryptophan (Trp) and methionine (Met)) have multiple encoding codons. Furthermore, tRNA wobble modifications fill in the gaps between codons and anticodons mismatch, allowing for non-cognate pairings at the third codon nucleotide that expands, and in some cases restricts, the decoding capacity of certain tRNA codons (2). This redundancy also allows for the dynamic finetuning of mRNA translation via altering codon optimality by dynamically changing tRNA modification levels to respond to external or internal cellular cues (3–7). For example, codon biased translation is increasingly being recognized as a driver for oncogenesis (5, 8) as well as a player in many other diseases (1). Thus, the classical view of codon usage and optimality is shifting from it being a static metric to a dynamic and adaptable one (3, 5, 9, 10).

Despite the ever-increasing interest in the mRNA translation level at the codon scale, and the presence of tools, such as Ribo-seq, that can provide such information, the dynamic nature of codon biased translation remains fundamentally incompletely understood. For example, why do certain tRNA modifications that occur in multiple tRNAs drive a pro-oncogenic codon biased translational program (5)? How can certain tRNA modifications, and their consequent impact on codon optimality, drive cellular stress response (3, 6, 7)? Are there global pro-oncogenic codon signatures that drive cancer progression? Previously, we have shown how tissue specific enrichment of tRNA modifications influence mRNA translation and codon bias in different mouse tissues (9). Other studies have highlighted the role of tRNA modifications in driving oncogenesis and other diseases (11, 12). However, a holistic view of codon biased translation is yet to be achieved.

In this study, we combined elements of evolutionary biology, computational analysis of coding sequences codon usage and bias, and analysis of multiple large proteomics datasets spanning human physiology, cancer cell lines, and human cancers, to reveal the evolutionary and functional relevance of codon usage bias. Importantly, our analysis validates the presence of distinct tissue signature of codon usage in humans, akin to what was observed in rodents (9) as well as the presence of a semi-global codon biased oncogenic signature that could be driven by several tRNA modifications.

## Results

### Analysis of mammalian Third nucleotide codon bias reveals contribution of amino acid content to GC3 drifts across species

To analyze codon bias in mammalian protein-coding transcriptomes we obtained coding sequences (CDS) for human (GENCODE v48), mouse (GENCODE vM37), and rat (Ensembl GRCr8). We selected one representative transcript per gene, opting to select the longest transcript for analysis. We employed 2 metrics for our analysis: GC3 scores and isoacceptors frequencies. GC3 scores were calculated by analyzing the codon sequences in a coding sequence, then calculating the ratio of codons ending in G or C (i.e. the third nucleotide is G or C) versus those ending in A or T (13). The score ranges from *zero* (no G/C ending codons) to 1 (all codons are G/C ending) [Figure 1A, see methods]. Isoacceptors frequencies were calculated as described previously (14, 15). Essentially, we compared the ratio of a given codon in a coding sequence to the sum of all synonymous codons of the same amino acid, yielding an isoacceptors score ranging from *zero* to 1 [Figure 1B, see methods]. We next created codon count tables and matrices for GC3 scores per gene and isoacceptors frequencies scores per codon per gene [Figure 1C]. To compare between the three species, we selected the common genes (≈15,000 protein coding genes) in all 3 data sets. We observed clear differences between human GC3 scores and those of the rat and mouse [Figure 1D]. While the GC3 density plots of the rat and mouse scores revealed a clear sharp peak around the median, the human GC3 scores were more plateaued with more genes having extreme GC3 scores. The median of the human GC3 scores remained like the other species (≈0.6). Spearman rank correlation analysis showed that while the correlation between the 3 species is high, it is higher between rat and mouse compared to their correlation with human GC3 scores [Figure 1E].

**Figure 1:**
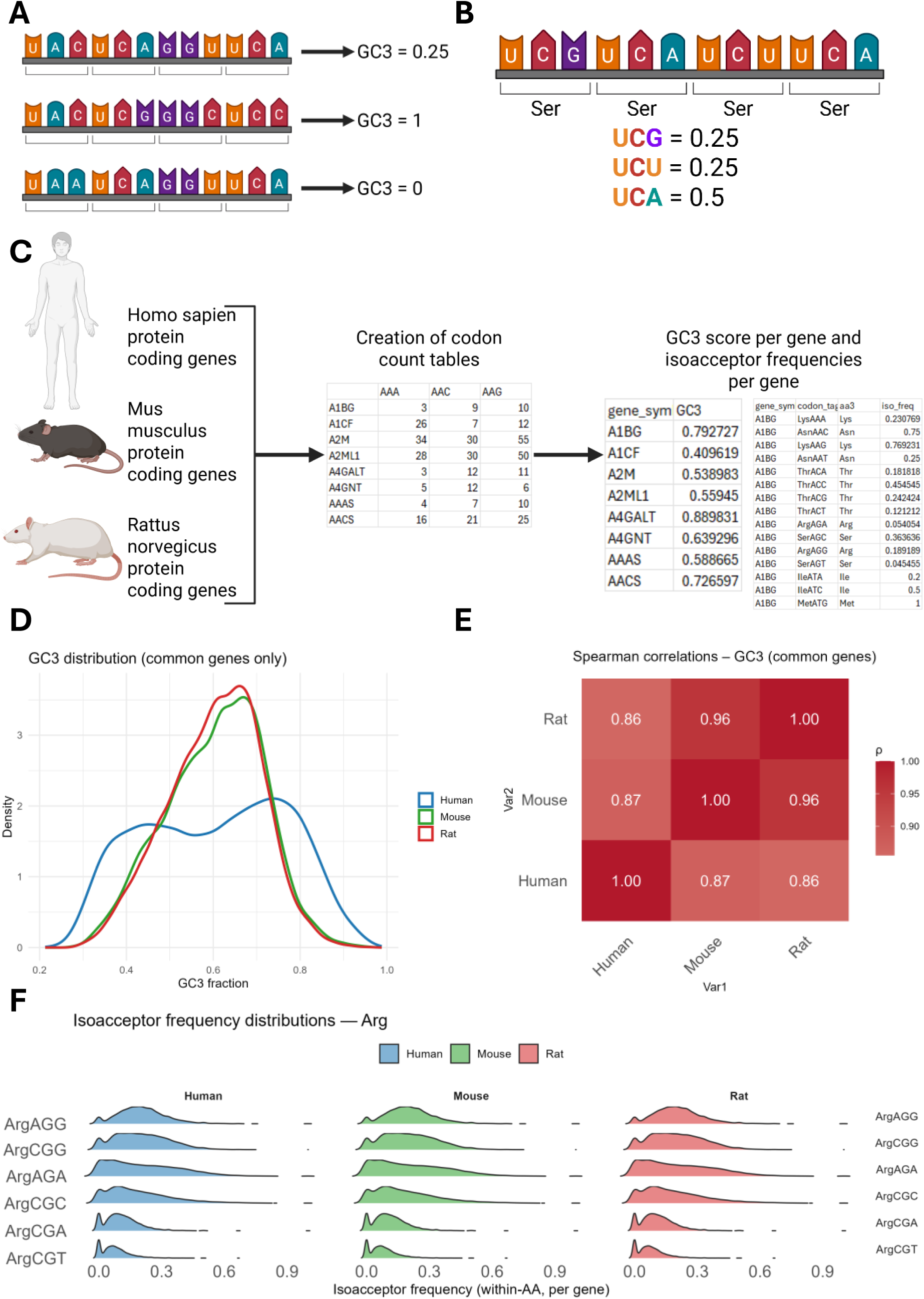
GC3 scores diverge between humans and rodents: **A:** An example of how GC3 score is calculated. **B:** An example of isoacceptors frequencies calculation. **C:** Design of this study. Coding sequences for human, mouse, and rat were retrieved and per gene codon count tables created. Using these count tables, per gene GC3 score and isoacceptors frequencies were calculated as explained in the methods section. **D:** Density plot of GC3 scores across the three species. **E:** Heatmap of Spearman’s rank correlation analysis of gene GC3 scores across the three species. **F:** Ridgeline analysis of isoacceptors codon frequencies of Arginine codons showing near identical values across all species. The same was evident for all other amino acid codons.

Despite these differences in GC3 scores between the three species, isoacceptors frequency scores were nearly identical for all codons [Figure 1F]. Thus, we could not attribute such differences to synonymous codon substitution. We argued that evolutionary changes in amino acid constitution of genes or gene length could explain such differences. First, we analyzed gene length differences between the three species. Globally, we observed statistically significant differences in coding sequence lengths across species [Supplementary figure 1A-B]. Next, we extracted the outlier genes based on their GC3 score differences between human and mouse and human and rat datasets by calculating ΔGC3 between human and each species separately (see methods). We selected arbitrarily the top 200 outlier genes for downstream analysis and observed that around 50% of those genes are shared between the human vs mouse and human vs rat GC3 analysis [Supplementary figure 1C]. Analysis of gene length of the outlier genes revealed statistically significant differences between rat outlier genes and other species but not between human and mouse [Supplementary figure 1D-E].

Next, we examined whether changes in amino acids composition between coding sequences of different species could explain the observed GC3 patterns, given that GC3 scores are sensitive to synonymous and non-synonymous codon substitution. To do so, we first collated the codons pertaining to each amino acid for each coding sequences to create amino acid count per gene.

Next, we normalized the counts by calculating the amino acid (AA) *z-*score for each gene. Plotting the z-scores for each amino acid across all genes and species revealed clear differences in some amino acids, such as Alanine, serine, and others [Supplementary figure 2A-B], while other amino acids had near identical internal *z-*score values across species [Supplementary figure 2C-D]. Sparse least square regression analysis (sPLS-DA) validated these differences in AA content of genes across the three species [Supplementary figure 2E]. Very important features analysis (VIP analysis) revealed that Aspartate, Arginine, and Alanine were the top determinants of clustering of genes across the three species [Supplementary figure 2F].

Next, we analyzed the outlier genes to examine whether their amino acid sequences could explain the changes in GC3 scores across species. First, we compared amino acid *z*-scores between outlier and non-outlier sets for each species. Next, we weighed observed differences by the intrinsic GC3 potential of each amino acid (i.e. the fraction of its synonymous codons that terminate in G/C). This weighting highlighted amino acids where differential usage is most likely to impact GC3 scores. T-test was used and adjusted *p*-values were obtained using the Benjamini– Hochberg method. We visualized the significant amino acids in our analysis as heatmaps of Δ*z* (outlier – background) values, ordered by GC3 potential [Supplementary figure 3]. Certain amino acids enrichment patterns were stable across species while others showed differences. For example, Aspartate had higher Δ*z* in human outliers than in other species while Ala had higher Δ*z* in rat outliers.

In summary, our results suggest evolutionary contributions of gene length and amino acid substitution in driving third nucleotide codon bias across mammalian genomes.

### Third nucleotide codon bias is linked to distinct molecular pathways

To understand how third nucleotide codon bias, reflected in GC3 scores, differs between species, we plotted the shared genes across the 3 species using their GC3 scores [Figure 2A]. Using K-mean clustering, we clustered the genes into 3 K-mean clusters, after multiple iterations. There were 2 large clusters, clusters C1 and C3, and a smaller cluster, cluster C2. GOBP ORA analysis showed that the genes in cluster C1, which had lower GC3 scores in human, enriched for pathways linked to mitotic and cell division, proliferation, and chromosomal functions [Figure 2B]. On the other hand, C2 cluster, which had higher GC3 score in human, was enriched for genes linked to synaptic and neuronal function and cell fate commitment [Figure 2C]. To evaluate whether these patterns could explain the global differences in GC3 scores across species (i.e. the presence of more extreme GC3 scores in human) we analyze the top and bottom genes by their GC3 scores (using a preset threshold: human: > 0.9 or < 0.3, mouse and rat: > 0.8 or < 0.3) using overrepresentation analysis (ORA) of gene ontology biological processes (GOBP). In all species, we observed that the AT-biased genes (i.e. lower GC3 scores) were enriched for pathways related to cell proliferation and mRNA processing [Figure 2D, showing only human data for privity]. while GC-biased genes were enriched for cell fate commitment and neuronal functions [Figure 2E, showing only human data for privity]. To validate the biased enrichment of GC3 (GC-biased) and AT3 (AT-biased) genes towards cell fate and neuronal pathways vs cell proliferation and mRNA processing, we, orthogonally, analyzed the human gene ontology biological processes (GOBP) pathways to evaluate their inherent bias. To do so, we focused on those pathways with ≥ 8 genes and calculated the average GC3 score per pathways using the mean GC3 score of the genes nested under the pathway. The median GC3 score of all GOBP terms hovered around 0.6 [Supplementary figure 4A]. GC3-rich GOBP terms were linked to differentiation and neuronal functions [Supplementary figure 4B], while AT3-rich GOBP terms were mostly linked to cell proliferation [Supplementary figure 4C]. We further conducted keyword search in the top 5% GC3- or AT3-rich terms and found that GC3-rich terms enriched for differentiation and signaling keywords while AT-rich terms enriched for RNA processing and chromosome related keywords [Supplementary figure 4D]. The same patterns were observed when we repeated the analysis using gene ontology cellular component (GOCC) and molecular functions (GOMF) terms [Data not shown]. To evaluate whether the same patterns are replicated across species, we repeated the analysis using mouse and rat GOBP terms. The same dichotomy between proliferation/RNA processing and differentiation/neuronal pathways was also observed [Data not shown]. Collectively, the differences between human and rat/mouse gene GC3 scores do not reflect a change in function, but rather, potential changes in protein structures, which could reflect evolutionary changes.

**Figure 2:**
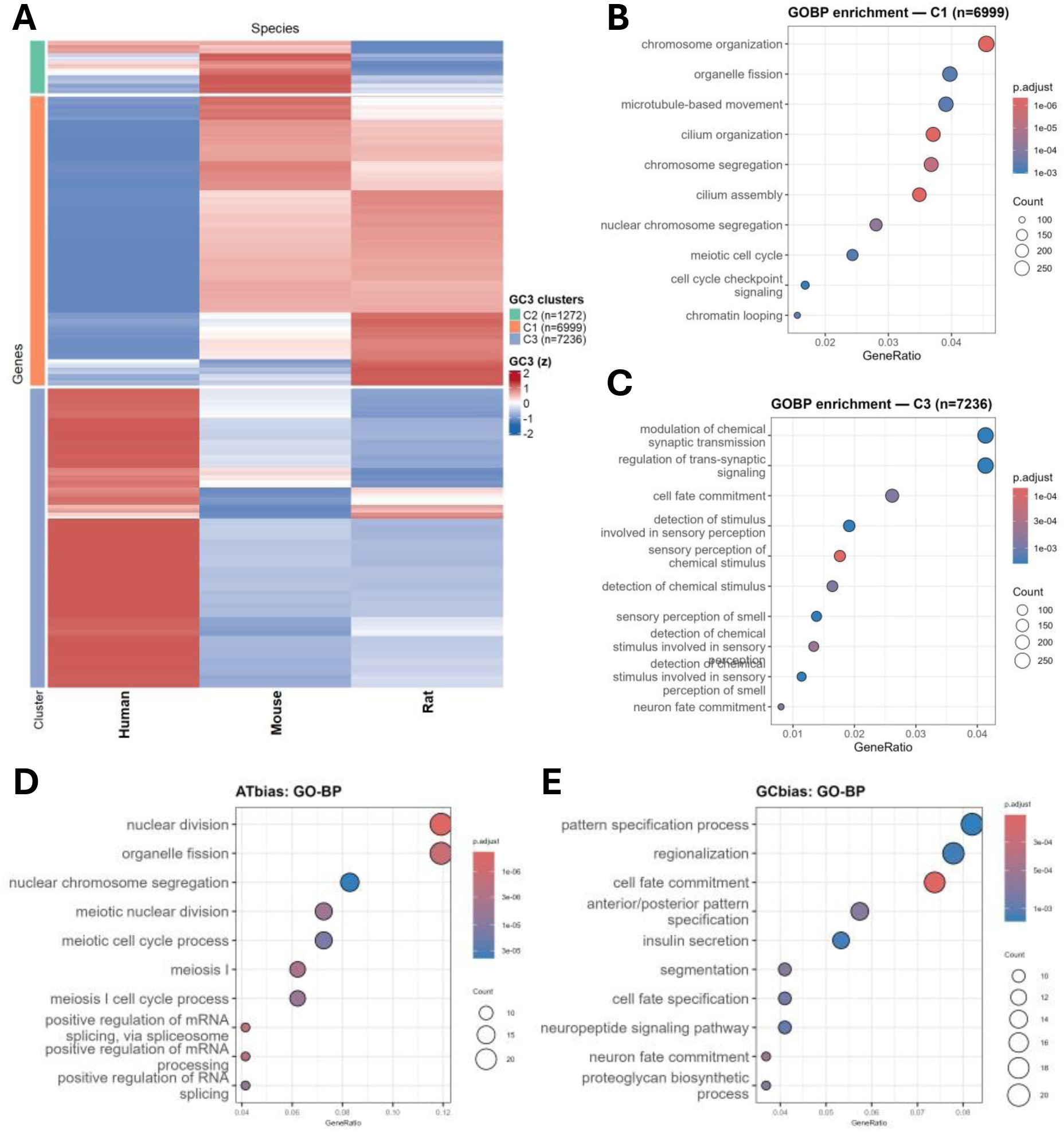
**A**: K-mean clustering of genes based on their GC3 across the three species examined. **B**: Overrepresentation analysis (ORA) of gene ontology biological processes (GOBP) of genes belonging to cluster 1 (C1) of the K-mean GC3 clusters. **C:** Overrepresentation analysis (ORA) of gene ontology biological processes (GOBP) of genes belonging to cluster 3 (C3) of the K-mean GC3 clusters. **D:** GOBP ORA analysis of the most A/T-ending biased genes. **E:** GOBP ORA analysis of the most G/C-ending biased genes.

To analyze this notion, we focused on the 2 main clusters showing changes in GC3 scores, C1 and C3 clusters [Figure 2A]. We extracted the sequence information of those genes and analyzed 3 parameters, GC content, gene length, and poly-A/P/Q tracts, which could reflect changes in amino acids sequences. Gene length in both clusters did not show great variations between human and rat/mouse genes [Figure 3A-B]. This negates the notion that gene length drives GC3 drifts, which we concluded when conducting genome-wide CDS length comparisons above. However, GC content showed clear statistically significant differences on Kolmogrov-Smirnoff statistical analysis between human and rat/mouse [Figure 3C-D]. These differences reflected the global trends in GC3 scores in the different clusters [Figure 2A], where C1 genes showed lower GC content in humans while C3 genes showed higher GC content in humans.

**Figure 3:**
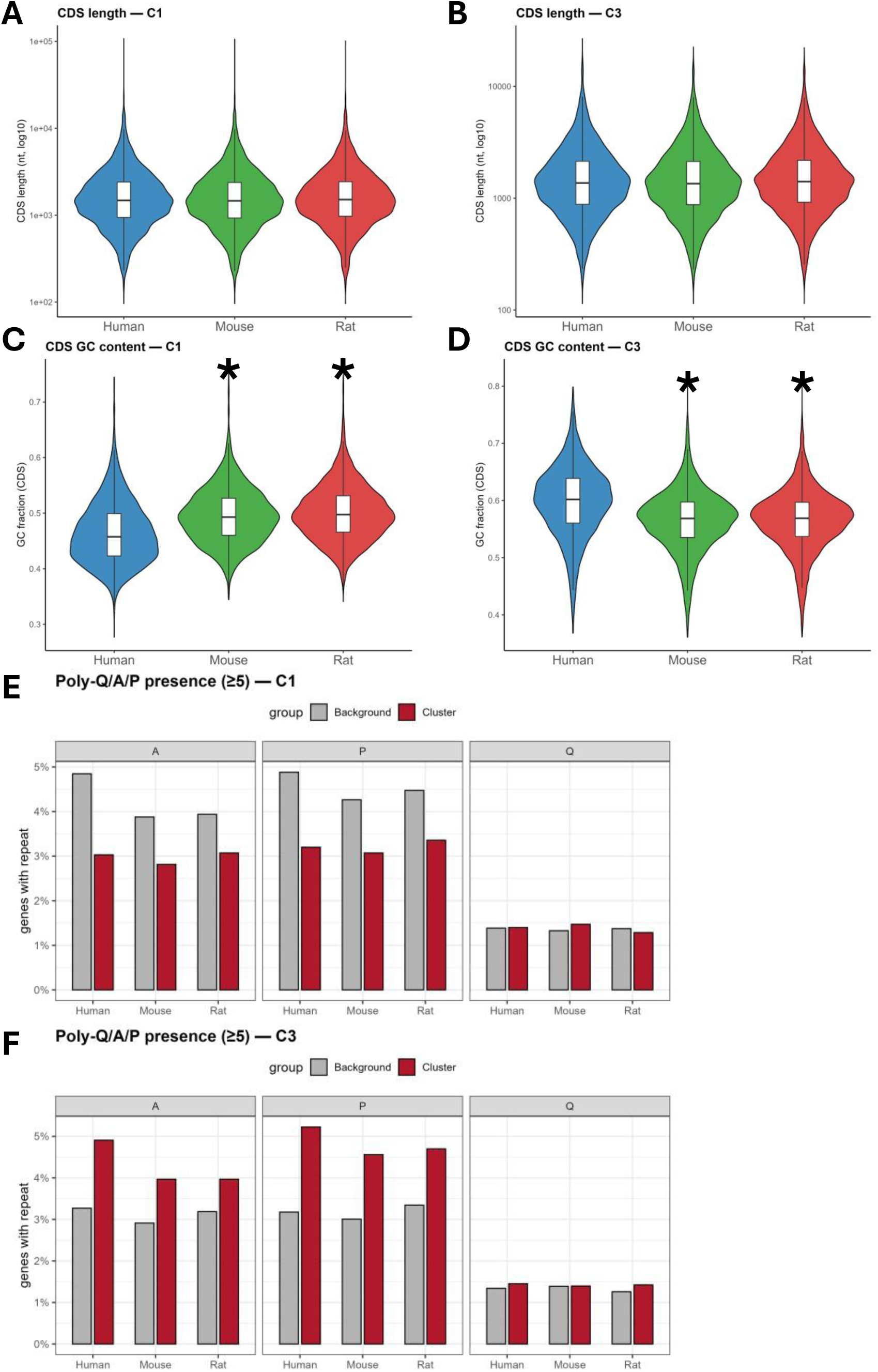
In-depth analysis of the 2 divergent GC3 clusters. **A-B:** Violin plots of CDS length of genes belonging to C1 or C3 across the three tested species. **C-D:** Violin plots of CDS GC-content of genes belonging to C1 or C3 across the three tested species. Asterisk indicates statistical significance compared to human CDS GC-content. **E-F:** Percentage of genes with Poly-A/P/Q repeats (minimum of 5 repeats) in C1 or C3 clusters compared to the background. After comparing it to the background, human set was compared to mouse/rat sets via Fisher’s exact test with FDR correction.

Next, we analyzed the genes in C1 and C3 clusters for the presence of poly-Alanine (poly-A), poly-Glutamine (poly-Q), and poly-Proline (poly-P) stretches, then compared poly-Q/A/P containing genes in each cluster with the whole genome, then statistically compared across species to observe if there are differences in the length of poly-Q/A/P tracts. We selected genes having a minimum of 5 poly-Q/A/P repeats for the analysis [Figure 3E-F]. Comparing the number of genes with poly-Q/A/P vs background and then across species using Fisher’s test with FDR correction revealed no significant differences between species in C1, but statistically significant differences in the numbers of poly-A containing genes between human and mouse/rat in C3 (FDR < 0.05). Additionally, we had two interesting observations. First, in both clusters, there were statistically significant differences in the number of genes in the cluster and in background in each species separately in poly-A/P tracts but not in poly-Q tracts. Secondly, we observed a reversal of the enrichment pattern between C1 and C3 clusters, where in C1, there were fewer genes containing poly-A/P tracts compared to the background and vice versa in C3 cluster.

Collectively, our data pointed out to a potential evolutionary GC-biased gene conversion (gBGC) due to recombination (16), which could explain the differences we observed in amino acid content of the coding sequences across species. To test such hypothesis, we initially calculated the delta-GC3 scores for each gene (ΔGC3), which we defined as: 𝛥𝐺𝐶3 = 𝐻𝑢𝑚𝑎𝑛 𝐺𝐶3 − 𝑀𝑒𝑎𝑛 𝐺𝐶3 𝑜𝑓 𝑀𝑜𝑢𝑠𝑒 & 𝑅𝑎𝑡. A positive ΔGC3 indicates a drift in human genes towards more GC-ending codons compared to rodents and vice versa. We selected genes with |ΔGC3| ≥ 0.2 to examine. The analysis yielded 133 human genes with GC3 gain (i.e. higher GC3 scores compared to the mouse orthologs) and 170 genes with GC3 loss (i.e. lower GC3 scores in humans). Analysis of gene length revealed statistical differences in both gain and loss gene sets between humans and mouse/rat orthologs [Figure 4A-B]. We selected the top gene by ΔGC3 score (SRSF8, ΔGC3 = 0.47) and the bottom gene (PARD6B, ΔGC3 = -0.38) for visualization [Figure 4C-D]. As seen in the 2 examples presented, recombination occurs leading to substitution of A/T with G/C and vice versa, thus leading to the GC3 drift observed.

**Figure 4:**
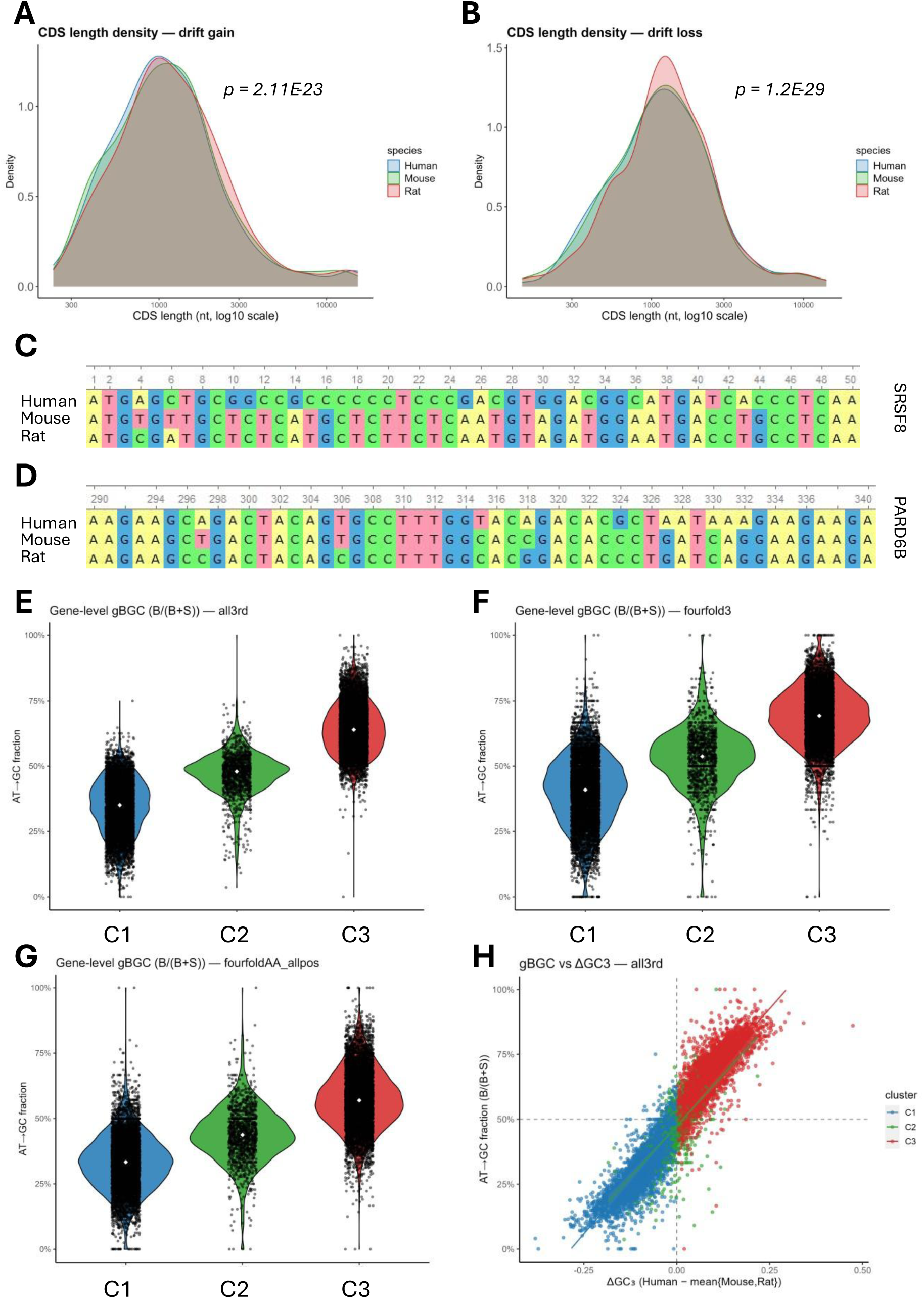
Synonymous and non-synonymous recombination events drive human GC-biased gene conversion. **A:** Density plot of CDS lengths of genes with delta-GC3 score gain in humans. P value indicates paired Wilcoxon statistical analysis between human and mean mouse/rat. **B:** Density plot of CDS lengths of genes with delta-GC3 score loss in humans (i.e. drift towards A/T-ending bias). P value indicates paired Wilcoxon statistical analysis between human and mean mouse/rat. **C:** Snippet of multiple alignment of SRSF8 CDS across species. **D:** PARD6B multiple alignment snippet across species. **E-G:** Volcano and swarm plots of GC-biased gene conversion (gBGC) analysis. The analysis was done for each of the 3 GC3 clusters in Figure 2A separately. Each dot represents a gene. Increase score indicates AT to GC conversion and vice versa. A score around 50% indicates neutrality or no significant conversion. **H:** Spearman’s rank correlation analysis between delta GC3 and gBGC scores.

Next, we analyzed gBGC by first creating per-gene codon alignment maps for all genes shared between the 3 species and identifying the rodent consensus sequence at the third codon position. We conducted the analysis in 3 passes to differentiate between synonymous and non-synonymous recombination. First, we analyzed the four-fold degenerate sites (i.e., amino acids where the change in the third codon nucleotide leads to synonymous change, for example, Ala which is encoded by GCU, GCC, GCA, and GCG). Next, we excluded the four-fold degenerate sites to identify recombination events leading to non-synonymous substitution (i.e. changing the amino acid sequence of the protein). Finally, we did a global pass to evaluate the global trends in recombination. We tallied the AT to GC or GC to AT recombination events at the third codon nucleotide in human genes compared to rodent genes to analyze recombination events that drive the drift in GC3 scores. We conducted our analysis on each of the three GC3 clusters [Figure 2A]. We observed in all events (synonymous and non-synonymous) [Figure 4E], in synonymous recombination events [Figure 4F], and in non-synonymous events [Figure 4G] that cluster 3 (C3) had higher AT to GC conversion events while cluster 1 (C1) had lower AT to GC conversion events (or higher GC to AT conversion events). Cluster 2 (C2) remained neutral with no significant drifts. These observations align with our clustering and the changes observed in GC3 scores between human and rodents. It is also notable that we observed a strong agreement between the ΔGC3 gene scores and their gBGC scores [Figure 4H], indicating that ΔGC3 can be an easy and quick way to screen for potential drifts due to recombination events.

In summary, we observed evolutionary drifts in third nucleotide codon composition between humans and rodents that lead to synonymous and non-synonymous codon substitution. However, the most AT-ending and GC-ending biased genes in all 3 species enriched for the same functional pathways, i.e., proliferation versus differentiation, indicating that these changes did not influence gene functions as expected in orthologs.

### Synonymous codons bias clusters genes to distinct groups

The translation of specific genes or gene sets in a given context is dependent on their GC3 scores (or 3^rd^ nucleotide bias) as well as isoacceptors frequencies (4, 6, 13, 14, 17). However, while our analysis revealed evolutionary changes in GC3 bias from rodents to humans due to non-synonymous and synonymous recombination, gene isoacceptors frequencies were globally conserved, apart from few outliers (data not shown). Indeed, there are differences between humans and species when examining specific outlier genes, as demonstrated by Davis et al (10), which makes sense if we consider synonymous recombination. Nonetheless, transcriptome-wide, we could not observe large drifts in gene isoacceptors frequencies as observed in the GC3 analysis. Davis et al (10) also showed that genes cluster by their isoacceptors frequencies into functional groups. We thus replicated their analysis by calculating per codon isoacceptors scores (ranging from 0 to 1, see methods and Figure 1B) for each gene using the human coding sequences (CDS). Here, we will focus on the human CDS for brevity. However, we replicated the same analysis for mouse and rat CDS and observed similar trends (Not shown).

Based on their per codon isoacceptors frequencies, we clustered the human genes into 10 k-mean clusters [Figure 5A]. Next, we conducted ORA GOBP analysis on each cluster to identify functional groups. Indeed, we observed unique functional groupings of clusters [Supplementary table 1]. To summarize our findings, we conducted keyword analysis on the GOBP enrichment of each cluster, identified unique keywords for each cluster, and used the top keywords to indicate functional grouping [Figure 5B]. We observed clear functional clustering based on isoacceptors codon frequencies, confirming Davis et al’s results (10). For example, Clusters 1 and 6 were enriched for immune-linked pathways, while Cluster 3 was enriched for neuronal and differentiation pathways. Cluster 4 was enriched for mitotic and cell division pathways. Clusters 5 and 8 were enriched for translation and RNA metabolism pathways. Cluster 7 was enriched for reproduction, embryogenesis, and development pathways. Cluster 10 was enriched for pathways linked to ion signaling, muscle contraction, and synaptic transmission. There were overlaps in the functional gene sets between clusters in addition to the presence of significant unique functional sets per cluster [Figure 5C]. We next evaluated what codons could be contributing to the clustering observed. To determine which codons were most informative for distinguishing between the k-means clusters, we quantified, for each codon, the proportion of its variance explained by cluster membership (between-cluster sum of squares / total sum of squares, BSS/SSₜ). This measure is equivalent to an ANOVA effect size (η²). Codons with higher BSS/SSₜ values therefore represent features that contribute strongly to cluster separation. Ranking codons by their BSS/SSₜ scores revealed a subset of highly informative codons [Figure 5D]. We next summarized the distribution of these top codons across clusters by calculating the average isoacceptors frequency for each cluster and centering it against the global background means. This yielded Δ isoFreq values, where positive values indicate codons enriched in a given cluster relative to the genome-wide baseline, and negative values indicate depletion [Figure 5E].Interestingly, we observed global patterns of A/T-ending vs G/C-ending codon enrichments across clusters in addition to codon specific enrichment patterns.

**Figure 5:**
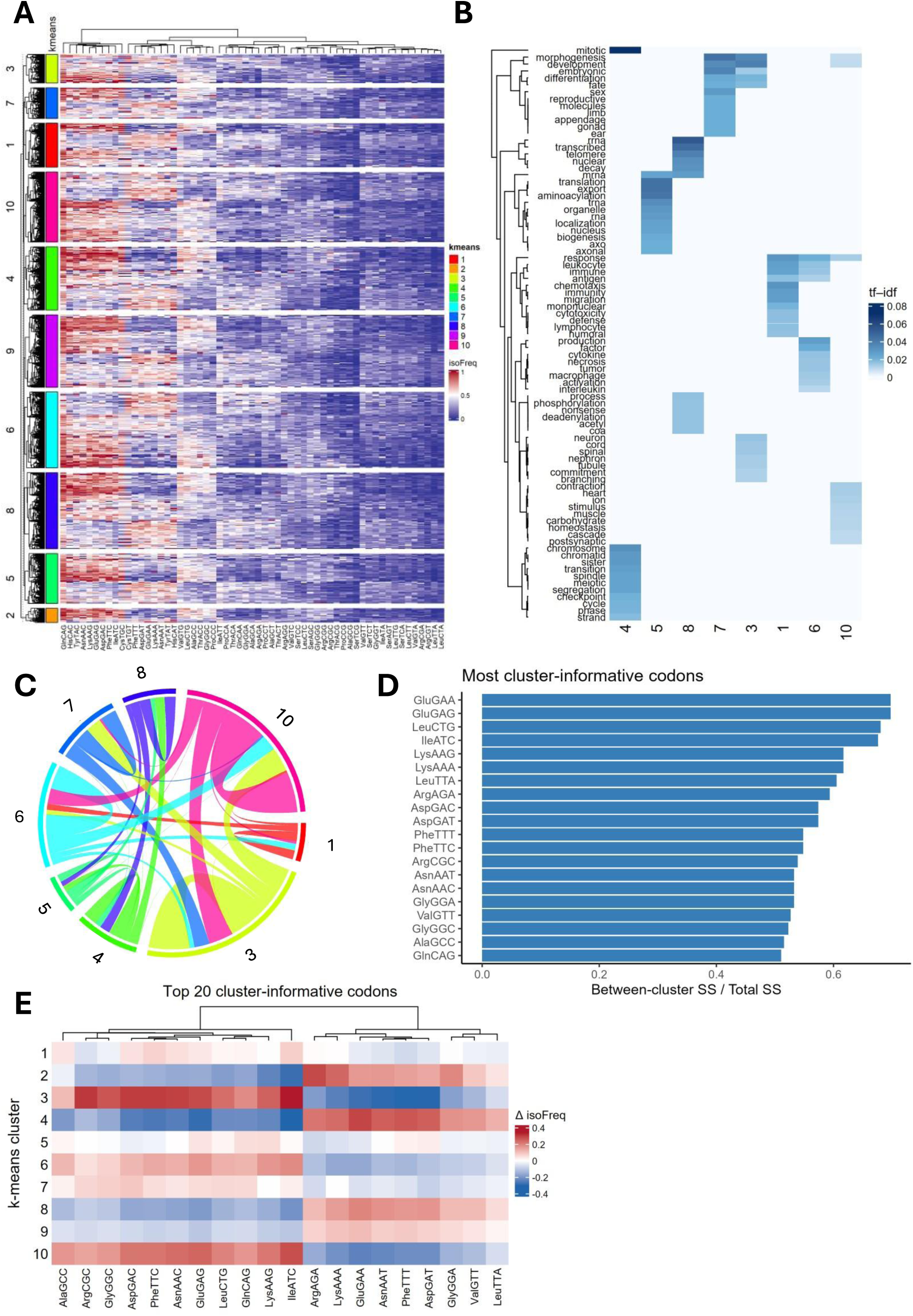
Isoacceptors frequencies cluster genes in distinct functional clusters. **A:** K-mean clustering of human genes by their isoacceptors frequencies. **B:** Heatmap of the top representative keywords of enriched GOBP terms in each cluster. **C:** Circos plot of unique and shared enriched GOBP terms in each cluster. **D:** Top 20 informative codons across clusters. Delta iso-freq indicate shift in mean iso freq of codon in that cluster compared to the background (all genes). **E:** Heatmap showing the Δ isoacceptors frequencies (isoFreq) of the top 20 informative codons in each cluster. Δ isoFreq was calculated by comparing the isoacceptors frequencies of the codon in a cluster versus the background.

Next, we wanted to examine whether GOBP pathways can also cluster based on their isoacceptors frequencies. To do so, we analyzed each GOBP term as a separate entity, focusing on those with ≥ 8 genes, calculated the isoacceptors frequencies for genes belonging to that term, and compared them to the genome average to generate T-stat values where |T-stat| ≥ 2 indicate statistical significance differences in codon isoacceptors frequencies (*p* < 0.05) compared to the background. We observed distinct pathway clusters as well as global A/T-ending vs G/C-ending clustering in GOBP terms [Supplementary figure 5A]. Furthermore, the top G/C-ending biased pathways were linked to development, neurogenesis, and cell signaling, while the top A/T-ending biased pathways were linked to cell cycle, proliferation, and RNA processing [Supplementary figure 5B-C]. These patterns are indeed mirroring the observations made during the GC3 analysis and reflect global patterns. However, such trends don’t clearly answer the question whether synonymous codons do regulate different processes or enrich in different pathways. This is of relevance when considering, for example, changes in codon decoding during cancer that derives oncogenesis (1, 5).

To explore such notion, we selected the top 5% genes by the isoacceptors frequencies of each codon and conducted GOBP ORA analysis to evaluate synonymous codons driven functional enrichment and compare between synonymous codons. Globally across all codons, we observed differential enrichment towards specific functional gene sets and pathways as well as clustering patterns that grouped certain codons together [Figure 6A]. We also observed differences in functional enrichment in genes enriched for synonymous codons. For example, Histidine (His) is encoded by 2 codons, CAC and CAU (CAT in the plots). His has only one anticodon, GUG, and thus requires queuosine (Q) tRNA modification to decode the CAU codon (18). Functional enrichment analysis revealed that His-CAC rich genes enrich for pathways linked to protein-DNA complexes, nucleosome assembly, and myeloid cells and megakaryocytes differentiation [Figure 6B]. On the other hand, His-CAU is linked to pathways related to immune response and mitochondrial respiratory complex assembly [Figure 6C]. Another expanded example is Leu, which is encoded by 6 codons. Leu-CTG is the most abundantly used codon in humans and rodents, while leu-CTA and leu-TTA are the rarest [Supplementary figure 6A]. We observed specific and distinct enrichment patterns across the 6 Leu codons [Supplementary figure 6B].

**Figure 6:**
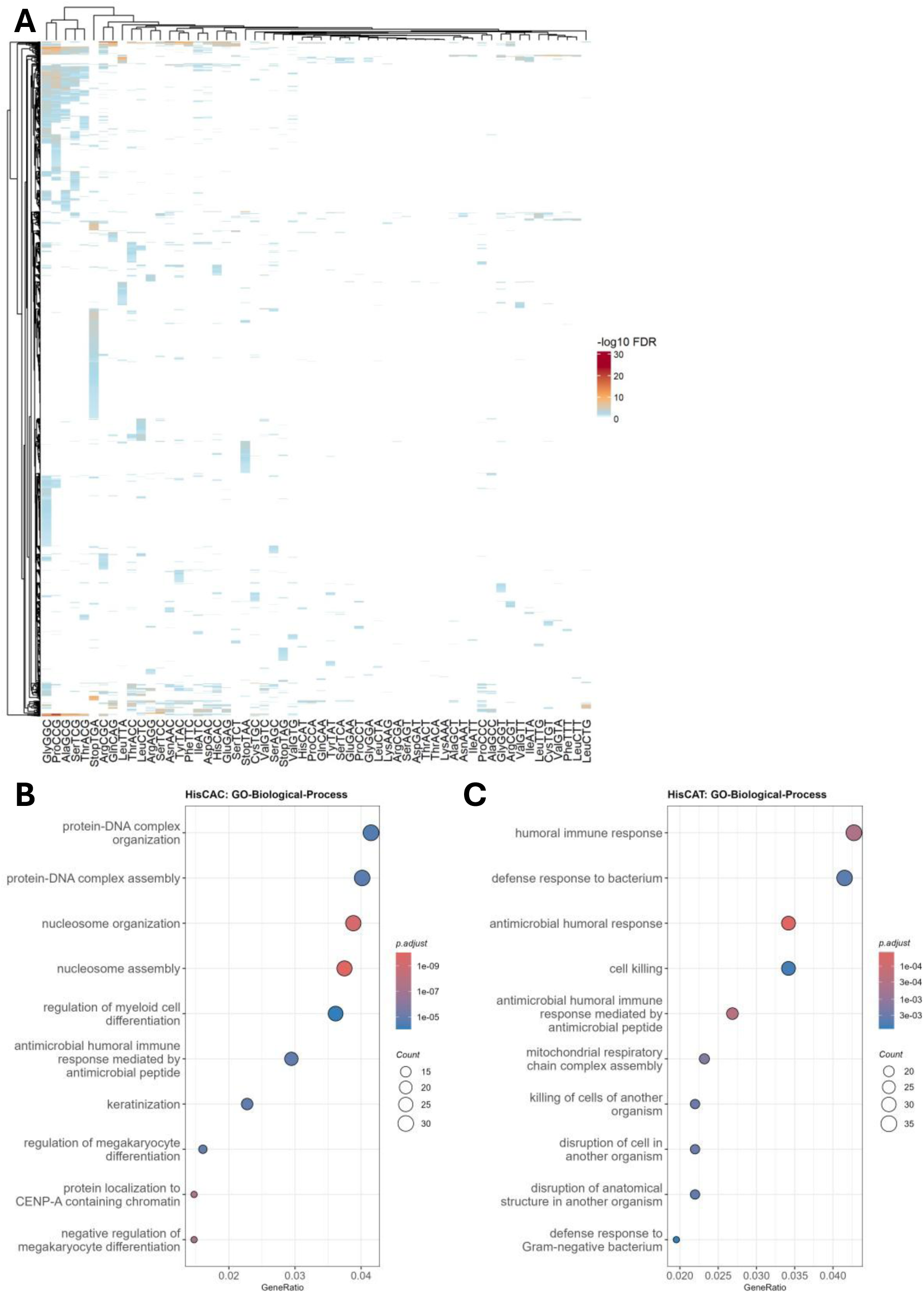
Analysis of synonymous codons influence on gene functional enrichment. **A:** Global heatmap of GOBP ORA analysis of genes enriched in different synonymous codons. After selecting the top 5% genes by their synonymous codon isoacceptors frequencies, GOBP ORA analysis was performed. The process was performed for all synonymous codons and the heatmap is a summary of all top GOBP terms enriched per codon. **B-C:** Example of enrichment analysis of synonymous codon pair. His-CAC or His-CAU biased genes were selected and GOBP ORA analysis performed.

Leu-CTG enriched for differentiation and developmental pathways as well as processes related to oxygen transport. Leu-CTC enriched mostly for ion transport related processes. Leu-TTG and Leu-CTT both enriched for pathways related to RNA processing and translation. Leu-CTA and Leu-TTA enriched for pathways linked to chromosomal division, vesicle localization, and cell cycle.

In summary, we show that codon isoacceptors frequencies or codon usage preference segregate genes into functional groups. Importantly, synonymous codon do enrich towards different processes/pathways, which is of great importance when considering, for example, that changes in tRNA modifications were shown to drive cancer oncogenesis or oxidative stress response by shifting translation towards specific codons and genes (5, 14, 19). Thus, our analysis clarifies why changes in tRNA modifications can shift cellular phenotypes even by changing the optimality of a synonymous codon pair.

### Gene codon content predicts impact of tRNA modifications

In recent years, our understanding of epitranscriptomic regulation of mRNA translation and disease pathologies has evolved drastically (1, 2, 5). Importantly, many tRNA modifications have been shown to promote cancer via codon biased translation (1, 5). Despite the increasing number of reports, it remains unclear why shifts in codon optimality are essential for cancer oncogenesis and progression. While the role of many tRNA modifications in cancer remain to be studied, there are few modifications that have been consistently linked to cancer aggressiveness. In particular, N7-methylguanosine (m^7^G) and N6-threonylcarbamoyladenosine (t^6^A) have been consistently reported to be linked to several cancers pathologies. To understand how these 2 modifications promote oncogenesis across multiple cancers, we analyzed mRNAs enriched for their codons. First, we examined t^6^A, which promotes the translation of ANN codons via codon-anticodon stacking (20). t^6^A is implicated in multiple cancers such as glioblastoma (19), hepatocellular carcinoma (21), and others. To understand the links between t^6^A and cancer, we created an ANN-index, whereas we counted, for each gene, the ratio of ANN codons compared to all codons (see methods). The median ANN-index across the genome was ≈0.3 [Supplementary figure 7A]. Next, we selected the top and bottom 10% genes by their ANN-index and conducted ORA GOBP analysis to examine their functional enrichment. Genes enriched in ANN codons enriched for mRNA translation and proliferation linked pathways [Supplementary figure 7B], while those poor in ANN codons enriched for differentiation linked GOBP terms [Supplementary figure 7C]. Given that this enrichment pattern mimics the proliferation versus differentiation pattern observed when we examined gene GC3 scores, we conducted Spearman’s rank correlation between gene ANN-index and GC3 score. We observed a significant negative correlation between these two indices (Spearman’s Rho = -0.632) [Supplementary figure 7D]. Thus, tRNA t^6^A levels increase in cancer shifts translation towards an A/T-ending biased pro-oncogenic program.

Next, we examined m^7^G, which is implicated in multiple cancers due to increased copy number of its writer METTL1 (5). We first created a consensus set of m^7^G decoded codons. We curated m^7^G modified tRNAs detected in 5 studies (11, 22–25) and selected those that appear in at least 3 studies as the consensus set. Next, we created the m^7^G-index by calculating the ratio of m^7^G decoded codons in a gene to all codons (See methods) [Supplementary figure 8A]. Genes rich in m^7^G codons enriched for pathways linked to mRNA translation, immunity, and metal ion stress response [Supplementary figure 8B], while those poor in m^7^G codons enriched for differentiation and development linked pathways [Supplementary figure 8C]. As with ANN-index, m^7^G-index negatively correlated with gene GC3 scores (Spearman’s Rho = -0.525) [Supplementary figure 8D], indicating that increased m^7^G levels in cancer cells drive an A/T-ending biased translational program.

In summary, this analysis reveals similar patterns of translational shifts in two important cancer-linked tRNA modifications that derive cancer proliferation, stemness, and immune evasion. In fact, we observed good agreement between gene ANN-index and m^7^G-index (Spearman’s Rho = 0.6) [Supplementary figure 8E], indicating similar oncogenic driving mechanisms at the codon level that globally shift translation towards A/T-ending codons.

### Analysis of the human proteome reveals distinct codon biases across tissues

Previously, it was shown that codon usage and bias influence mRNA translation and gene expression at the tissue level (13, 17). Importantly, in mouse tissues, most abundantly translated mRNAs were A/T-ending biased except for the brain, which was more G/C-ending biased (17). Understanding these patterns is important from the standpoint of developing more specific gene and mRNA therapeutics via codon reengineering to achieve better protein expression in target tissues/cells and reduce expression in off-target sites (10, 17). To that end, we retrieved an atlas of 32 human tissues proteomes and transcriptomes from the GTEx database (26) and analyzed the codon patterns across tissues. Spearman’s rank correlation showed a modest correlation between RNA and protein expression across tissues, with a global Spearman’s Rho of 0.299 and a range from as low as 0.145 (Breast mammary tissue) to 0.471 in the cerebellum [Figure 7A]. Next, for each tissue, we selected the top 10% expressed genes/proteins and analyzed their GC3 scores and isoacceptors frequencies. At the RNA level, all tissues appeared to be G/C-ending biased with similar isoacceptors frequencies patterns across the board [Figure 7B]. However, at the protein level, we observed clear clustering of G/C-ending versus A/T-ending biased tissues [Figure 7C], further confirming the transcriptome-proteome mismatch. Importantly, the clustering mimicked what was observed previously in mice using Ribo-seq (17) where the brain was G/C-ending biased while most tissues were A/T-ending biased. In human proteomes, tissues from the nervous system, arteries, skin, lungs, and spleen were G/C biased. On the other hand, Liver, heart, muscles, and other internal organs were mostly A/T-ending biased. The testis was observed to be the most A/T-ending biased tissue as well as the tissue with the strongest difference between RNA and protein GC3 scores [Figure 7D]. This dichotomy fits the narrative of A/T-ending bias being linked to proliferation and G/C-ending bias being linked to neuronal function and differentiation given our knowledge of tissue physiology. To further clarify such notions, we conducted ORA GOBP analysis on the top 10% expressed proteins in each tissue [Supplementary figure 9A]. We observed general clustering trends mimicking What was observed at the protein isoacceptors frequencies analysis. Further, GOBP ORA analysis of the most A/T-biased tissue (the testis) [Supplementary figure 9B], and the one of the most G/C-biased tissues (The cerebellum) [Supplementary figure 9C] recapitulated the proliferation versus differentiation/neuronal function bias observed at the level of global A/T-ending versus G/C-ending bias. Examining the GC3 scores of the top 20 GOBP terms enriched in each tissue revealed the clear G/C versus A/T-ending preferences for different tissue [Supplementary figure 10A-B].

**Figure 7:**
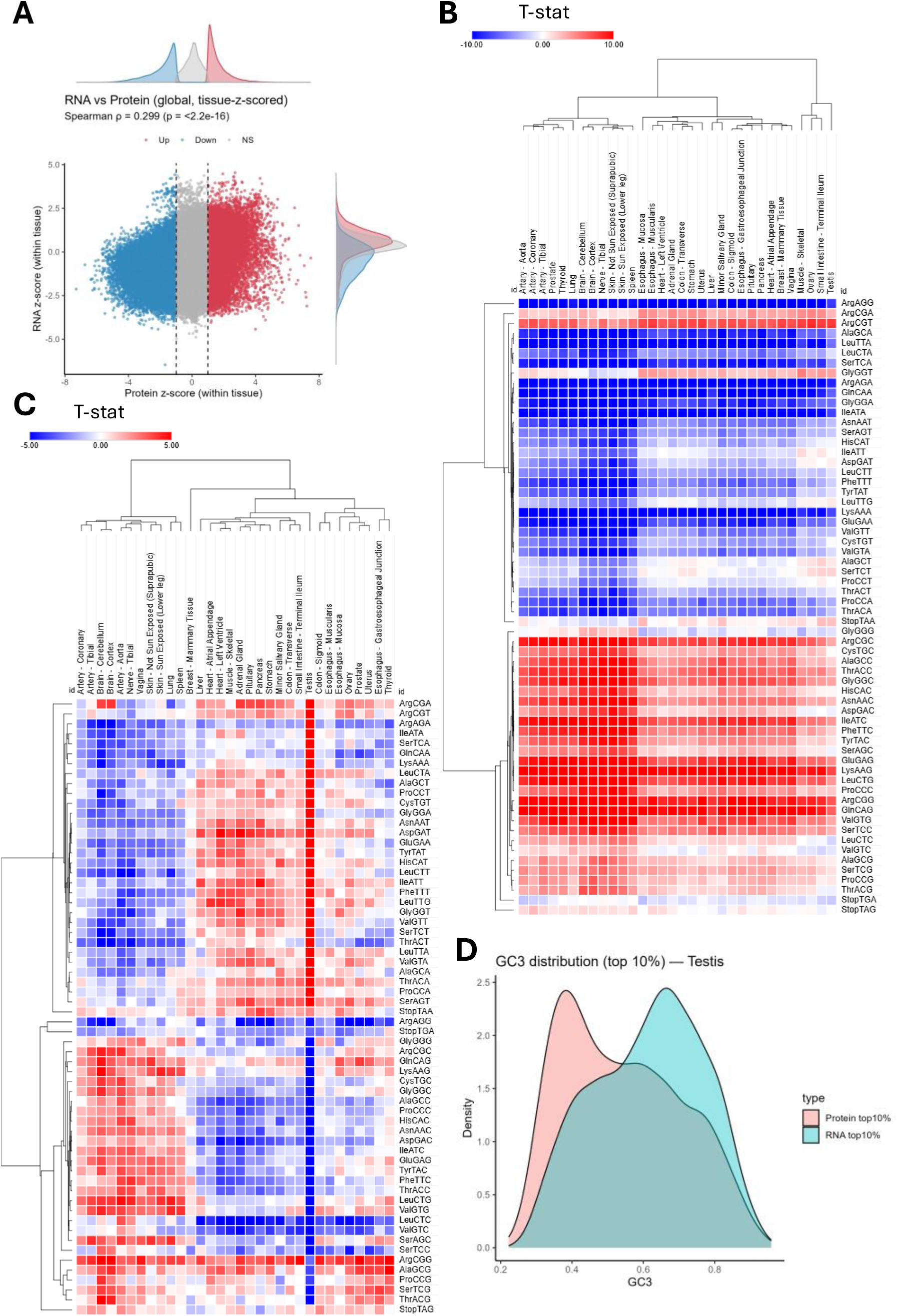
Analysis of human tissues proteomes and transcriptomes. **A:** Spearman’s rank correlation analysis of RNA and protein expression across all tissues. **B:** heatmap of isoacceptors codon frequencies analysis of top 10% expressed RNAs in human tissues presented as T-stat of tissue expressed genes versus the genome as background. **C:** heatmap of isoacceptors codon frequencies analysis of top 10% expressed proteins in human tissues presented as T-stat of tissue expressed genes versus the genome as background. **D:** GC3 density plots of the top 10% expressed mRNAs and proteins in testis.

In summary, we observe tissue specific codon patterns that clusters human tissues based on their synonymous codon biases in agreement with previous computational analysis (13) and model organisms based analysis (17, 27). Importantly, we observed strong differences between RNA and protein expression as well as codon bias when analyzed at the RNA or the protein levels, indicating a strong regulatory influence at the level of mRNA translation regulating codon selection and bias that impacts the proteome (4, 14). Our observations indicate that tissue specific codon-biased translation are fine-tuned to tissue’s physiological functions.

### Analysis of a cancer cell line atlas reveals heterogeneity in codon usage across cancer cell lines

The importance of understanding codon decoding extends from evolutionary biology to understanding diseases and developing therapeutics. As shown above, certain cancer-linked tRNA modifications drive oncogenesis via codon biased mRNA translation. Given the ongoing interest in understanding epitranscriptomics, tRNA modifications, and codon-biased mRNA translation in driving cancer and as potential therapeutic avenue (5, 8), we extended our analysis to 2600 cancer cell lines analyzed in cell model passports database (https://cellmodelpassports.sanger.ac.uk/) (28). We first conducted quality control (QC) analysis on the whole dataset, which showed variations in the numbers of models (i.e., cell lines) related to different tissues [Supplementary figure 11A] and the number of representative tissue status (e.g. tumor vs metastasis) in the database [Supplementary figure 11B]. Importantly, the number of detected genes and proteins showed acceptable variability and normal distribution across the database [Supplementary figure 11C-D]. Spearman’s rank correlation revealed nearly no correlation between RNA and protein expression (Spearman’s Rho = 0.181 globally across all models). GC3 scores based on proteomics data showed relatively tight distribution across all models with no major variations [Figure 8B]. Next, we calculated the isoacceptors T-stat of the top 10% expressed proteins in the 948 models that remained after filtering and plotted them using K-mean and hierarchical clustering [Figure 8C]. We observed the presence of several major clusters, with one cluster being significantly G/C-ending biased (K-mean cluster 1) while another cluster was A/T-ending biased (K-mean cluster 2) [Figure 8C]. Examining the K-mean clusters profiles revealed overlaps between models originating from different tissues, indicating inter-tissue heterogeneity in terms of isoacceptors frequencies [Supplementary figure 12A]. To examine this notion, we analyzed models by tissues of origins. We observed significant inter-tissue heterogeneity across all tissues and models. For example, in cell models from central nervous system (CNS) and from those from hematopoietic and lymphoid tissues, we observed distinct clusters based on isoacceptors codon bias [Supplementary figure 12B-C]. Thus, K-mean clustering cannot be attributed only to specific tissue of origin of the model but rather indicates a more complex figure. Notwithstanding these observations, we noted the presence of patterns of enrichment of different tissue origins, cancer types, tissue status, and growth properties of the models across the clusters [Supplementary figure 13]. For example, cells growing as suspension and cells derived from metastatic tumors were more represented in cluster 1 [Supplementary figure 13C-D]. More granular analysis of cancer type revealed specific clustering. For example, breast carcinoma was more represented in cluster 2 while B-Cell Non-Hodgkin’s lymphoma was more represented in cluster 1 [Supplementary figure 13B].

**Figure 8:**
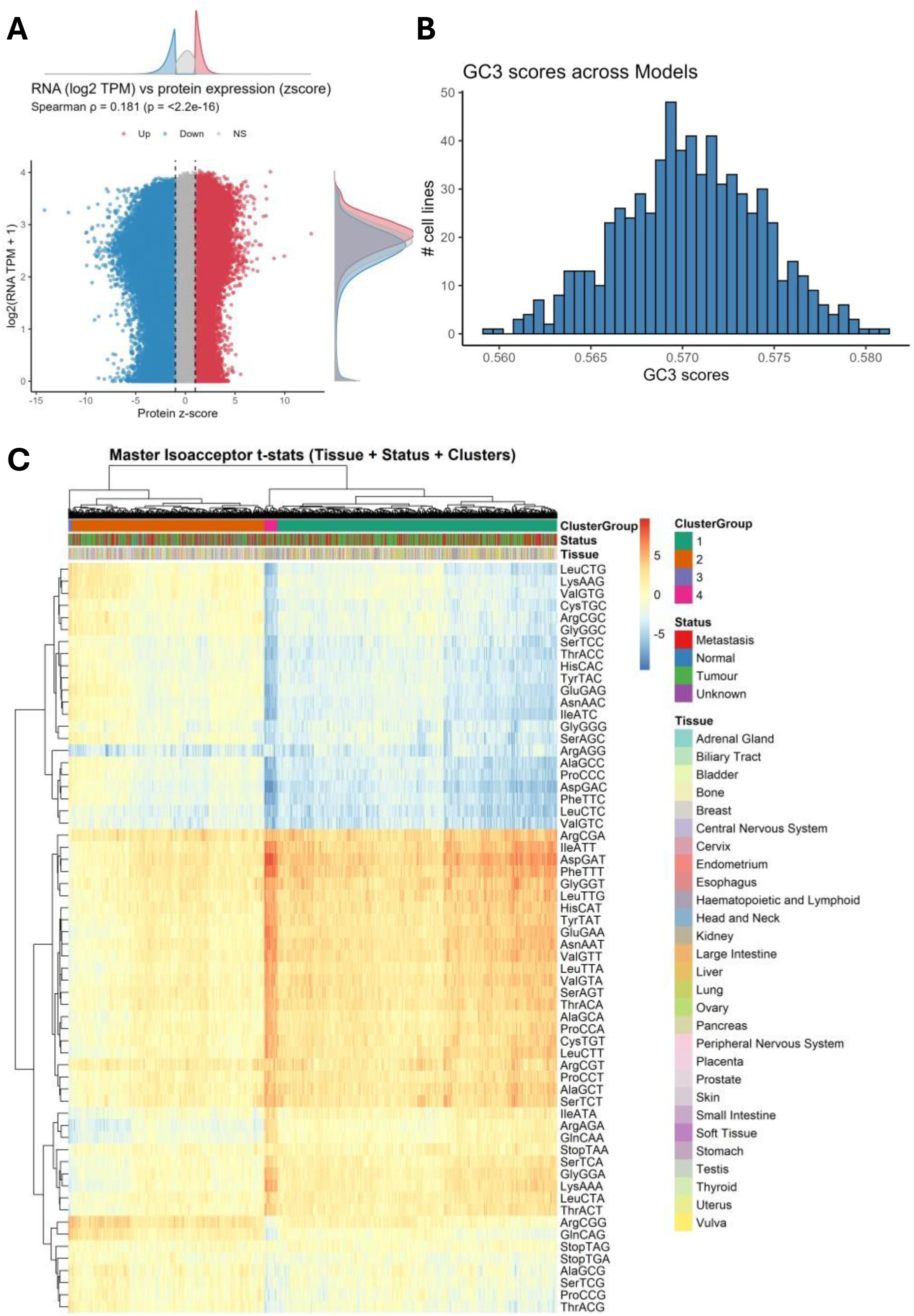
Analysis of cancer cell line atlas reveals heterogeneity in codon usage. **A:** Spearman’s rank correlation analysis of RNA and protein expression across all examined models (i.e., cell lines). **B:** Histogram of top 10% expressed proteins GC3 scores across all examined models. **C:** Heatmap with K-mean clustering of isoacceptors frequencies T-stat of the top 10% expressed proteins in all tested models.

We wanted to understand whether this heterogeneity in cancer cell lines codon usage is an inherent cancer feature due to patient heterogeneity or is due to culturing conditions. To do so, we retrieved human cancer samples proteomics from a recently published dataset (29). The dataset comprised of 999 tumor samples representing 22 cancer types. We retrieved the preprocessed protein intensities per sample, normalized the protein expression, selected the top 10% expressed proteins per sample, and then analyzed the isoacceptors frequencies for each sample to evaluate patient to patient heterogeneity in codon usage across cancers. We observe that within each of the examined tumors, isoacceptors codon frequencies were largely homogeneous, with few variations in certain codons across samples [Supplementary figure 14]. Thus, we can conclude that the heterogeneity of the codon usage observed in cancer cell lines is a feature of the cell lines and not due to the parent tumor itself. However, this should be validated by comparing cell lines and their tumor of origin using the same methodologies. Unfortunately, such datasets are not available in the literature.

In summary, we observed heterogeneous codon usage patterns across the many cell lines used in cancer research, which was not observed when we examined human cancer samples proteomes. This heterogeneity raises questions regarding the replicability of findings related to mRNA translation dynamics, for example due to tRNA modification changes or other epitranscriptomic features, across different cell lines. Thus, there is a need to expand studies investigating translational changes in cancer cells to include different cell lines with different codon usage dynamics to ensure the robustness of observations and relevance to clinical cancer features.

### Analysis of human cancer proteomes reveal global A/T-biased translational shifts

Given that we hypothesized that a global A/T-ending codon biased translation, driven, for example, by certain tRNA modifications such as t^6^A or m^7^G, could be a hallmark of cancer, we wanted to test such notion in human cancer specimens. To that end, we retrieved information of different cancers proteomes from CancerProteome database (http://bio-bigdata.hrbmu.edu.cn/CancerProteome/index.jsp) (30). The database includes an analysis of different public databases and provides information of differential expressed proteins in cancer samples versus normal tissues, which allowed us to examine A/T-ending codon biased translational shifts in cancer. Using this information, we extracted the upregulated and downregulated proteins in 21 cancer types provided by the database and analyzed their GC3 scores and isoacceptors frequencies. Globally, we observed that upregulated proteins were more A/T-ending biased while the downregulated proteins were more G/C-ending biased based on the GC3 scores of the differentially expressed proteins [Supplementary figure 15A]. However, certain cancers were G/C-ending biased such as acute myeloid leukemia [Supplementary figure 15B]. Isoacceptors frequencies analysis offered a more granular view of codon usage bias, revealed specific codon preferences across cancer types, and confirmed the global cancer A/T-ending codon shifts, with the presence of few exceptions [Supplementary figure 15C-D]. Thus, we can argue that global A/T-ending codon bias, shifting translation towards a proliferation program, could be defined as a hallmark of many cancers.

## Discussion

In this work, we utilized 2 important metrics to understand how codon biased translation can shape mRNA translation and link protein output to functionality. GC3 scores give information on global codon bias in genes, and whether the genes are A/T-ending or G/C-ending codon biased (13). Isoacceptors codon frequencies, or synonymous codon bias, gives a more granular view on the preference of synonymous codons in a given coding sequence (31). We observed that GC3 scores evolutionary drift from rodents to human genomes due to synonymous and non-synonymous gene recombination, validating previous observations (16), without alteration in gene functional enrichment. On the other hand, these drifts did not lead to major alterations in gene isoacceptors codon frequencies, apart from few outliers (10). In both analyses, we observed functional stratification of genes based on their codon content. A/T-ending versus G/C-ending codon bias (derived from GC3 scores) revealed a proliferation versus differentiation/neuronal signature in both gene-level and pathway-level analyses (32). Synonymous codon usage analysis revealed a more granular view of functional gene enrichment and stratifications. Importantly, we show how different enrichment of synonymous codons could drive divergent translational programs, as shown in the provided examples of His and Leu codons.

We also show for the first time how tRNA modification indices, such as the ANN-index and m^7^G-index used in this work, can explain how tRNA modifications are linked to oncogenic programs (5). Such indices indicate that a global A/T-ending biased translation program could be a hallmark of cancer. A finding we observed in many cancers when we analyzed human cancer proteomes compared to their tissues of origin (30). While in most cancers, we observed the expected A/T-ending biased translational/proteome program, there were exceptions. Thus, future works profiling tRNA expressions and modifications in different cancers will be essential in understanding these patterns globally across all cancers and in a more detailed way in different cancer types and subtypes. When examining cancer cell lines, we observed stark diversity in codon usage across cell lines derived from the same tissue and cancer origins. This figure was not observable when we examined human cancer sample proteomes. Thus, we can deduce that this heterogeneity represents a feature of the cancer cell lines due to culturing conditions or other changes occurring during their development. Nonetheless, this heterogeneity raises important questions and concerns regarding the interpretation of results from different studies using different cell lines and calls for the application of some measure for reporting findings from cancer cell lines. For example, studies focusing on translation dysregulation in cancer should employ different cell lines with different codon usage programs to evaluate whether the reported observations are context specific to a cell line or global to a cancer type/subtype.

One of the interesting applications of understanding codon usage bias in healthcare is the codon reengineering of mRNA/gene constructs to achieve better protein production in target cells/tissues while reducing the aberrant off-protein production. We previously demonstrated this principle in mice tissues using *in vivo* adenovirus mediated gene delivery of mutant EGFP constructs (9). Davis et al (10) also showed the same principal *in vitro*. Thus, a complete understanding of codon usage at the tissue, cell, and cell state is essential for future biotechnological applications. We show here in human tissues that different tissues have different codon usage signatures, replicating, with some differences, our observations in mice tissues (9). Importantly, these patterns were not observable when we examined mRNA expression datasets of the same tissues, highlighting the importance of translational regulation in dictating codon usage and proteome output. We further observed distinct codon usage signatures when we examined human cancer and cancer cell lines proteomes. Nonetheless, it is important to note that current approaches in studying cancer translation or proteomics are done in-bulk. Thus, the true cancer cell signatures might be diluted or altered due to the presence of other cells in the samples such as immune cells. However, our computational approach indicates that a global A/T-ending codon bias signature, which we observed in most cancers’ proteomes, is a cancer codon signature. Previous works also alluded to the presence of such signatures in cancers. For example, Rapino et al (33) showed that wobble uridine tRNA modifications drive the translation of proteins enriched in AAA, GAA, and CAA codons in BRAF(V600E)-expressing melanoma cells. The upregulation of these proteins was linked to resistance to chemotherapy in melanoma.

While the analysis here focused on cancer datasets, as they are the most curated and high quality available, it is important to note that the applications extend beyond the cancer field. Translational deregulation and the consequent proteostasis dysregulation are known to occur in many conditions such as aging (34, 35), diabetes (12, 36), and neurodegenerative diseases (37) to name a few. Understanding global and specific codon patterns in disease conditions could, in theory, allow for the design of more robust gene/mRNA therapeutics that can achieve better results in target cells.

It is important to highlight that our analysis and conclusions regarding cancer codon usage shifts are limited by the quality of the datasets available. Technical variations or low-quality samples could indeed skew the analysis. While we selected datasets from studies that we believe are of high quality, more orthogonal analysis and validation is required to fully comprehend the influence of codon usage and bias on cancer initiation, progression, and outcomes.

## Conclusions

We show here that codon usage and bias segregate genes into functional groups and that codon biased translation is finetuned to meet the needs of tissues and cells. Importantly, we report an A/T-biased cancer codon signature that could be further explored to optimize gene and mRNA therapeutics designed to target cancer cells. We believe that more research on mRNA translation in cancer, and importantly creation of epitranscriptomic and translatomic datasets of cancers at the tissue and cell level will be essential in understanding cancer pathophysiology and in designing novel therapeutics.

## Methods

### Data sources and preprocessing

Protein coding sequences (CDS) for human and mouse transcriptomes were retrieved from Gencode (v48 for human and vM37 for mouse). Rat coding sequences were retrieved from Ensemble (GRCr8). CDS were harmonized and the longest CDS per gene was selected for downstream analysis. Codon count tables were generated by coRdon package in *R*.

The human tissue transcriptomes and proteomes were retrieved from a previously published article (26). RNA-seq expression matrices (log₂-transformed TPM) and proteomics relative abundance matrices were imported, and gene identifiers were mapped from Ensembl IDs to HGNC gene symbols to homogenize all datasets and files.

The cancer cell lines analysis was conducted using cell models passport database (https://cellmodelpassports.sanger.ac.uk/) (28). The RNA-seq (2025-02-17) and proteomics (2025-02-11) datasets were downloaded as csv or tsv files and directly imported into *R* after harmonization for analysis. Metadata of cell lines annotation (2025-04-23) and gene identifiers (2024-12-12) were also downloaded and used in our analysis.

Human cancers with matched normal tissues proteomics were retrieved from CancerProteome database (30). Differentially expressed proteins (FC and FDR values) in 21 cancer types were downloaded from the database and used for the analysis.

Per-tumor sample proteomics were retrieved from a published study (29). Per-sample protein intensities were downloaded from the supplementary data of the study and used for downstream analysis after z score normalization.

### GC3 score analysis

The GC content at the third codon position (GC₃) was calculated per gene as:

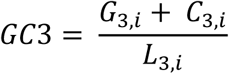

where 𝐺_3,𝑖_ + 𝐶_3,𝑖_ is the sum of codons having guanine or cytosine nucleotides at position 3 for gene *i*, and 𝐿_3,𝑖_ the total number of codons in the gene.

To compare across species, mean rodent GC₃ was defined as:

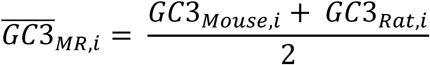

The human-specific drift was quantified by:

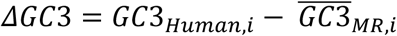

|ΔGC3| ≥ 0.2 indicated drift genes. Drift genes were selected for downstream analysis.

### ANN-index

The ANN-index was calculated by this equation:

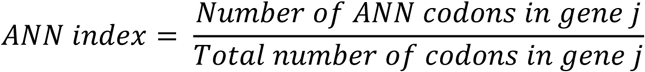

### m^7^G index

m^7^G-index was calculated first by identifying consensus m^7^G modified tRNAs, defined as those detected in at least 3 studies from 5 studies that profiled m^7^G at the isoacceptors level (11, 22–25). The m^7^G-index was calculated as such:

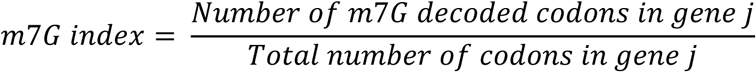

### Isoacceptors frequencies analysis

For each gene, codon usage was decomposed into frequencies of tRNA isoacceptors (synonymous codons decoding the same amino acid). Isoacceptors frequencies of a gene were calculated by:

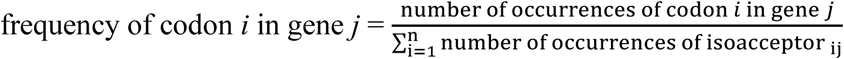

This yielded a score from 0 to 1.

### T-statistics for isoacceptors frequencies vs background

The T-statistic describing the isoacceptors codon frequencies of a list of selected genes was calculated by one-sample T-statistic against the genome-wide background:

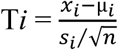

Where T*i* is the codon isoacceptors frequencies, x_i_ is the mean frequency for codon *i* in the sample, µ_𝑖_ refers to the mean frequency for codon *i* in the background (genome-wide), s_i_ refers to the genome-wide standard deviation of isoacceptors frequency for codon *i*, and n refers to the sample size (i.e., number of foreground genes or selected genes). The resulting T-stat value from the analysis indicates direction of enrichment (positive versus negative) and statistical significance (|T-stat| ≥ 2 indicates *p* ≤ 0.05)

### Poly-amino acid repeat detection (Poly-Q/A/P analysis)

After K-mean clustering of human, mouse, and rat genes by their GC3 scores, we identified the genes in each cluster and used them for downstream analysis. Protein sequences were translated from CDS and scanned for runs of glutamine (Q), alanine (A), and proline (P). For each gene, the longest uninterrupted repeat length was extracted. Genes were classified as “present” for a poly-run if the maximal run length was greater than or equal to a minimum threshold (*m = 5*).

Statistical testing included:

- **Presence/absence analysis:** pairwise Fisher’s exact tests with Benjamin-Hochberg multiple test correction were applied across species for each cluster to identify presence or absence of Poly-Q/A/P containing genes in each GC3 cluster versus the background as well as between species.
- **Repeat length comparison:** Kruskal–Wallis tests were used to assess differences in run lengths across species. Pairwise Wilcoxon rank-sum tests with Benjamini–Hochberg correction were conducted for species pairs.

### GC-biased gene conversion (gBGC) analysis

To analyze recombination events at the third codon nucleotide that could drive drifts in GC3 scores, we first created codon-level multiple sequence alignments per gene for all genes detected in all 3 species (15,348 genes). Next, we identified consensus nucleotide in the rodent (mouse and rat) coding sequences and compared the human to it. If we observed a change from A/T to G/C in humans we counted it as B (biased towards GC), if we observed a change from G/C to A/T we counted it as S. Next, we calculated the gBGC score as such:

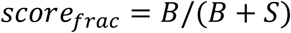

A higher *score_frac_* Indicates recombination events at the third codon nucleotide substituting A/T with G/C and leading to higher GC3 score and vice versa. A *score_frac_* around 50% indicates no significant drifting. The analysis was conducted in 3 passes, first, we analyzed all codons for synonymous and non-synonymous recombination events (all3rd), then we analyzed only four-fold degenerate codons (codons whose change in the third nucleotide leads to no change in encoded amino acid) for synonymous recombination events (fourfold3), and lastly we excluded the four-fold degenerate codons to analyze non-synonymous recombination events (fourfoldAA_allpos).

### Amino Acid z-Score Analysis

For each protein-coding gene and species, amino acid usage frequencies were converted into z-scores to normalize for background distribution. First, using the codon count tables, we translated each codon to its amino acid and created an amino acid count table. Next, we calculated the amino acid z-score per gene as such:

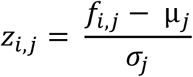

Where 𝑧_𝑖,𝑗_ is the standardized usage of amino acid *i* in gene *j*. 𝑓_𝑖,𝑗_ is the amino acid raw count, µ_𝑗_ is the mean amino acid counts in the gene *j,* and 𝜎_𝑗_ is the standard deviation of all amino acid counts in the gene.

Amino acid z-scores were calculated for all genes across species, then we selected the genes present in the 3 species for downstream analysis by Sparse Partial Least Squares Discriminant Analysis (sPLS-DA) and variable importance in projection (VIP) Using mixOmics package (38).

To identify amino acid changes in outlier genes (i.e. those that showed differences between species in the sPLS-DA analysis), we first calculated the amino acid z-score shift in outliers using this equation:

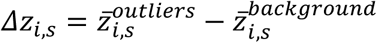

Where 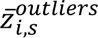 = mean z-score of amino acid *i* in outlier gene of species *s*, and 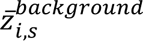 = mean z-score in non-outlier genes. Statistical significance was assessed with a two-sample t-test for each (species × amino acid) pair, and false discovery rates were controlled by the Benjamini– Hochberg method.

Because amino acids differ in their capacity to modulate GC3 (due to codon degeneracy), we quantified each amino acid’s **GC3 potential** as the proportion of its synonymous codons ending in G or C:

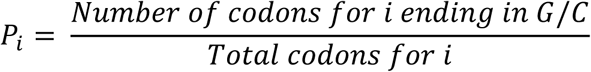

We then calculated a GC3-weighted shift:

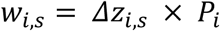

This weighting prioritizes amino acids whose codon structure allows stronger influence on GC3 variation. Heatmaps of 𝛥𝑧_𝑖,𝑠_ were generated, with amino acids ordered by GC3 potential (highest to lowest). Rows corresponded to amino acids and columns to species. A diverging color scale (blue = negative shift, red = positive shift, white = no change) was centered at zero. This representation highlights amino acids where outlier genes are enriched or depleted relative to background, and links those shifts to their potential impact on GC3 composition.

### Isoacceptors clustering and codon-level feature importance

for the isoacceptors-based clustering analysis, we used per-gene isoacceptors frequencies as calculated above. Genes were clustered using k-means (k = 10, 100 random starts, seed fixed for reproducibility) on the gene × codon isoacceptors frequency matrix after centering and scaling.

To identify codons contributing most strongly to cluster separation, we calculated, for each codon, the ratio of between-cluster sum of squares (BSS) to total sum of squares (SSₜ) across genes. This statistic, equivalent to ANOVA η², quantifies the proportion of codon variance explained by cluster membership. Codons with high BSS/SSₜ values were considered highly cluster informative. Codons were ranked accordingly, and the top 20 codons were selected for visualization.

For each cluster, centroid isoacceptors frequencies were computed as the average frequency of each codon among genes assigned to that cluster. To highlight enrichment or depletion, centroid values were mean-centered against the genome-wide codon averages, producing ΔisoFreq values. Positive ΔisoFreq values indicate codons enriched in a given cluster relative to the background, whereas negative values indicate depletion. These deviations were visualized in heatmaps with rows corresponding to clusters and columns to codons.

To assign functional interpretation to clusters, we performed over-representation analysis (ORA) of Gene Ontology Biological Processes (GOBP) using the clusterProfiler R package. Enrichment results were corrected for multiple testing using the Benjamini–Hochberg method and only significant enrichment (FDR < 0.05) were selected.

To provide an overview across clusters, we performed keyword analysis of significant GO-BP and KEGG terms. Enrichment descriptions were tokenized into keywords, stop words removed, and term frequencies weighted by term frequency–inverse document frequency (TF–IDF). The top-ranked keywords per cluster were used to summarize biological functions associated with codon-usage clusters.

### Isoacceptors codon-specific ORA enrichment

To investigate whether synonymous codons enrich for distinct functional categories, we ranked genes by isoacceptors frequency for each codon. For each codon, the top 5% of genes (highest isoacceptors frequency) were selected, and ORA GOBP analysis performed.

We compared enrichment results across synonymous codons of the same amino acid to assess functional divergence. Representative examples included Histidine (CAC vs CAT) and Leucine codons, where enrichment patterns were distinct despite encoding the same amino acid. Enrichment was summarized as dot plots of significant GO-BP categories per codon.

### Isoacceptors analysis of GOBP terms

Here, we analyzed each GOBP term as an independent entity, keeping our analysis to terms with ≥ 8 genes. For each GO-BP term, we computed the isoacceptors frequencies per codon across its member genes, then compared this to the genome-wide average to calculate T-stat values (see above). Deviations were converted into T-statistics (see below). The resulting codon × pathway matrix of T-statistics was clustered and visualized as a heatmap (ComplexHeatmap), allowing detection of global AT-ending versus GC-ending codon biases across pathways.

### Keyword analysis of GOBP enriched terms in isoacceptors clustering analysis

For each cluster GOBP ORA analysis, we tokenized the titles of the GOBP terms and removed English stop words and tokens with fewer than 2 alphabetic characters. Next, we calculated token raw counts (*n*) per cluster then calculated the relative token frequency:

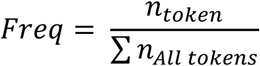

Next, we calculated the tf-idf scoring using this equation:

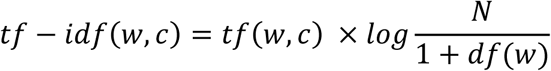

where *c* is cluster, *N* the number of clusters, *tf* is the term frequency, *idf* is the inverse document frequency, or how unique the token is across all clusters, and *df(w)* the number of clusters containing word *w*. For each cluster we report the **top 15** words by tf–idf (ties not allowed). Next, we visualized the enriched keywords per cluster as a heatmap with clustering based on shared keywords between clusters.

### GC3 analysis of GOBP terms

As with the isoacceptors frequencies analysis of GOBP terms, we selected terms with ≥ 8 genes. For each term, we calculated the mean GC3 score of genes nested under the term to generate per-term GC3 score.

### Keyword enrichment analysis of GO terms

To identify and summarize differences between biological processes associated with codon bias, we performed a keyword enrichment analysis on Gene Ontology Biological Process (GO-BP) terms stratified by GC3 content. GO-BP terms were first ranked by their mean GC3 index across annotated genes. The top and bottom 5% of terms were extracted and treated as “high-GC3” and “low-GC3” groups, respectively. Term names were tokenized into individual words, lowercased, and cleaned by removing standard English stop words and terms shorter than three characters.

Word frequencies were then calculated separately for the high- and low-GC3 groups. To quantify relative enrichment, we computed log₂ fold-changes in frequency between the two groups, adding a pseudo-count to avoid division by zero. The resulting “keyness” scores highlighted keywords disproportionately associated with high-versus low-GC₃ pathways. For visualization, the top 10 positively and negatively enriched words were plotted as a bar chart, enabling direct comparison of thematic trends between GC3-rich and GC3-poor biological processes.

## Acknowledgment

The authors report no conflict of interest nor any ethical adherences regarding this work.

## Author contributions

**S.R.:** Study conception and design. Funding acquisition. Formal analysis and interpretation. Manuscript writing. Administration. **K.N.:** Critically revised the manuscript. Funding.

## Data availability

No new data was generated for this manuscript. All data analyzed are available via public repositories or via the cited publications.

## Code availability

No new software was created for this study. All analyses were done in *R* studio. The scripts used can be requested from the corresponding author.

