## Supplementary figures for "Third-nucleotide codon bias and synonymous codon bias define functional translational programs that shape human tissue and cancer proteomes"

**Supplementary figure 1**: **A:** Violin plot of CDS length in the three tested species. **B:** Pairwise Kolmogorov-Smirnoff statistical analysis of CDS length. **C:** Venn diagram of the top 200 outlier genes based on GC3 score differences between species. Pairwise differences in GC3 values were calculated between human–mouse and human–rat. For each contrast, the top 200 genes with the largest absolute differences were designated as outlier sets. The overlap between outlier sets was quantified, and unique and shared subsets were extracted for downstream analyses. **D:** Violin plot of CDS length between outlier genes across species. **E:** Pairwise Kolmogorov-Smirnoff statistical analysis of CDS length of outlier genes.

A

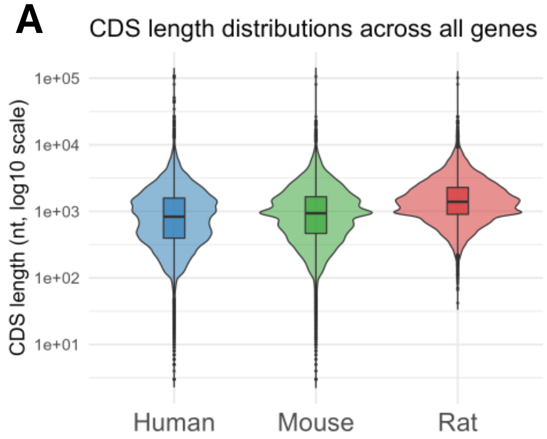


B

| contrast | D | *P* value |
|---|---|---|
| Human vs Mouse | 0.063087 | 5.98E-127 |
| Human vs Rat | 0.282962 | 1.00E-300 |
| Mouse vs Rat | 0.229651 | 1.00E-300 |

C

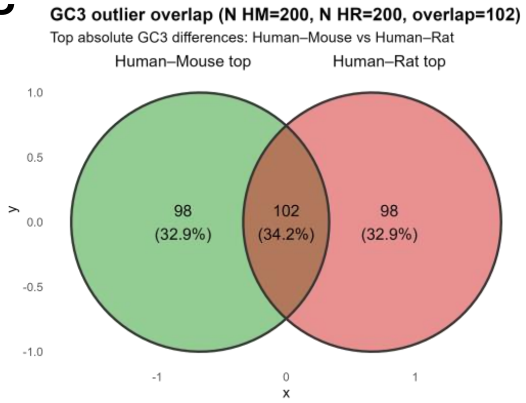


D

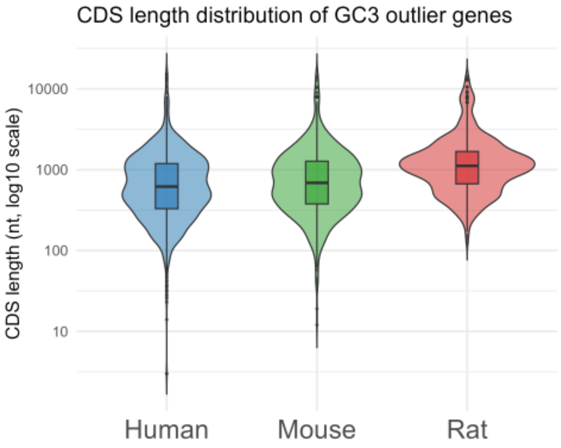


E

| contrast | D | *P* value |
|---|---|---|
| Human vs Mouse | 0.057962 | 0.057485 |
| Human vs Rat | 0.296628 | 4.89E-38 |
| Mouse vs Rat | 0.253889 | 1.21E-21 |

**Supplementary figure 2: A:-D:** Density plot of different amino acid z scores across species showing examples of divergent amino acid z scores **(A-B)** and similar scores (**C-D)**. **E:** Sparse partial least square regression discriminant analysis (sPLS-DA) of amino acid z scores across species. Each dot represents one gene. Each gene is colored by its species of origin. **F:** Variable Importance in Projection (VIP) analysis of sPLS-DA amino acid z scores analysis revealing the amino acids contributing to the clustering of genes across species.

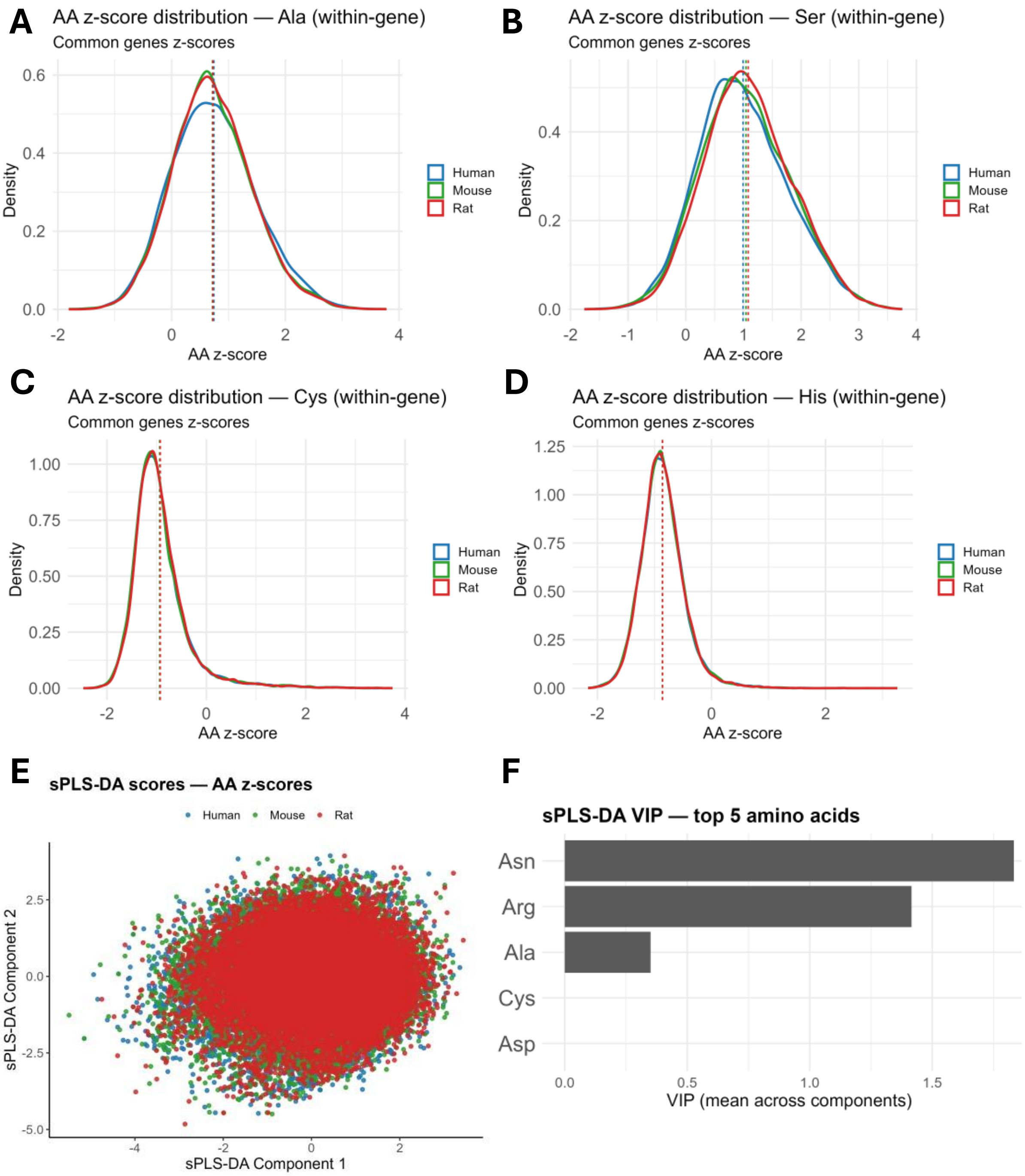
A
AA z-score distribution — Ala (within-gene)
Common genes z-scores
Density
AA z-score
Human
Mouse
Rat
B
AA z-score distribution — Ser (within-gene)
Common genes z-scores
Density
AA z-score
Human
Mouse
Rat
C
AA z-score distribution — Cys (within-gene)
Common genes z-scores
Density
AA z-score
Human
Mouse
Rat
D
AA z-score distribution — His (within-gene)
Common genes z-scores
Density
AA z-score
Human
Mouse
Rat
E
sPLS-DA scores — AA z-scores
Human
Mouse
Rat
sPLS-DA Component 2
sPLS-DA Component 1
F
sPLS-DA VIP — top 5 amino acids
Asn
Arg
Ala
Cys
Asp
VIP (mean across components)

**Supplementary figure 3: Analysis of amino acid contribution to variations in GC3 scores across species**. For each species and amino acid, *t*-tests were performed to compare amino acid *z*-scores between outlier and non-outlier sets. To evaluate the extent to which specific amino acids could contribute to GC3 variation, we weighted observed differences by the intrinsic GC3 potential of each amino acid (i.e. the fraction of its synonymous codons that terminate in G/C). This weighting highlight amino acids where differential usage is most likely to impact GC3 scores. Adjusted *p*-values were obtained using the Benjamini–Hochberg method. Results were visualized as heatmaps of $\Delta z$ (outlier – background) values, ordered by GC3 potential, allowing rapid identification of amino acids contributing to species-specific codon usage differences.

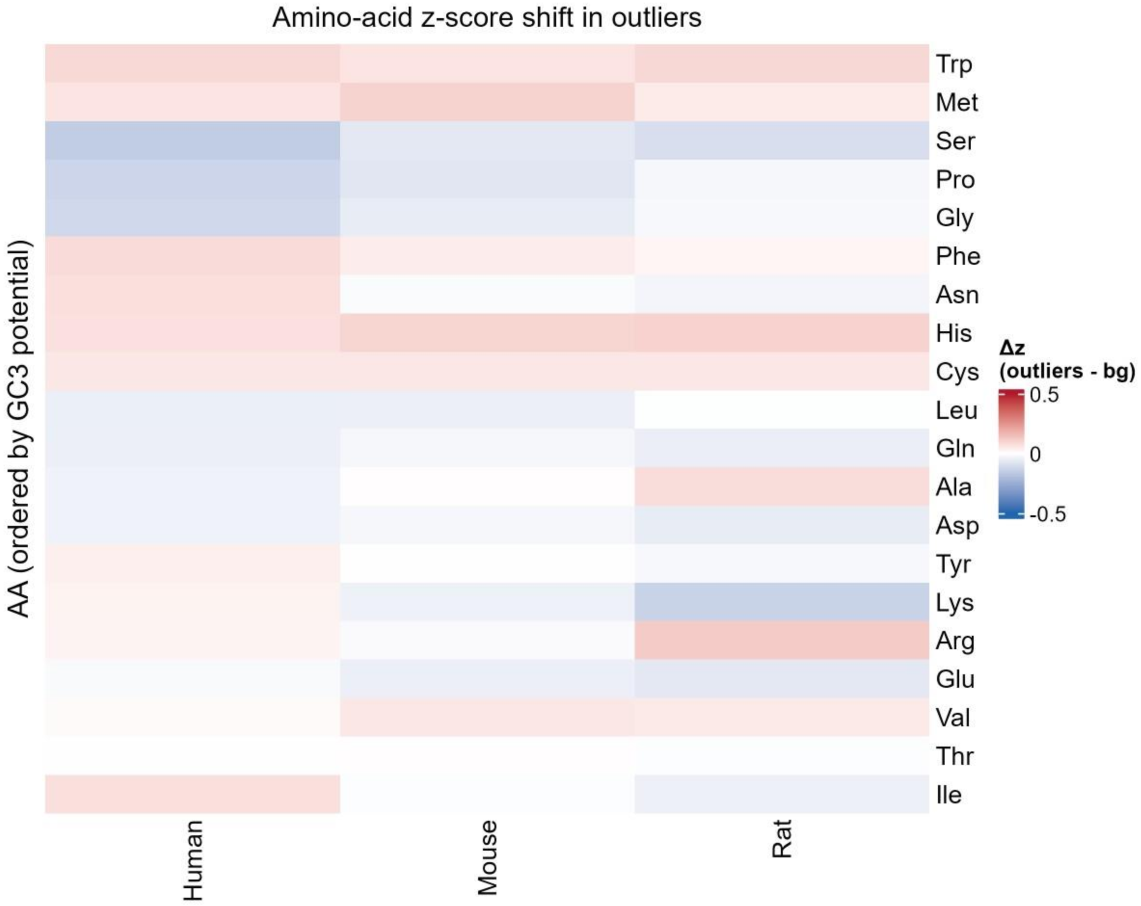

Amino-acid z-score shift in outliers
AA (ordered by GC3 potential)
Trp
Met
Ser
Pro
Gly
Phe
Asn
His
Cys
Leu
Gln
Ala
Asp
Tyr
Lys
Arg
Glu
Val
Thr
Ile
Human
Mouse
Rat
Δz
(outliers - bg)
0.5
0
-0.5

**Supplementary figure 4: Analysis of GOBP terms GC3 scores**. **A:** Density plot of all mean GC3 scores of GOBP terms with >= 8 genes. **B:** Top 10 GC3-rich GOBP terms. **C:** Top 10 AT3-rich terms. **D:** Bar plot showing keyword enrichment analysis of the top and bottom 5% GOBP terms by their GC3 scores.

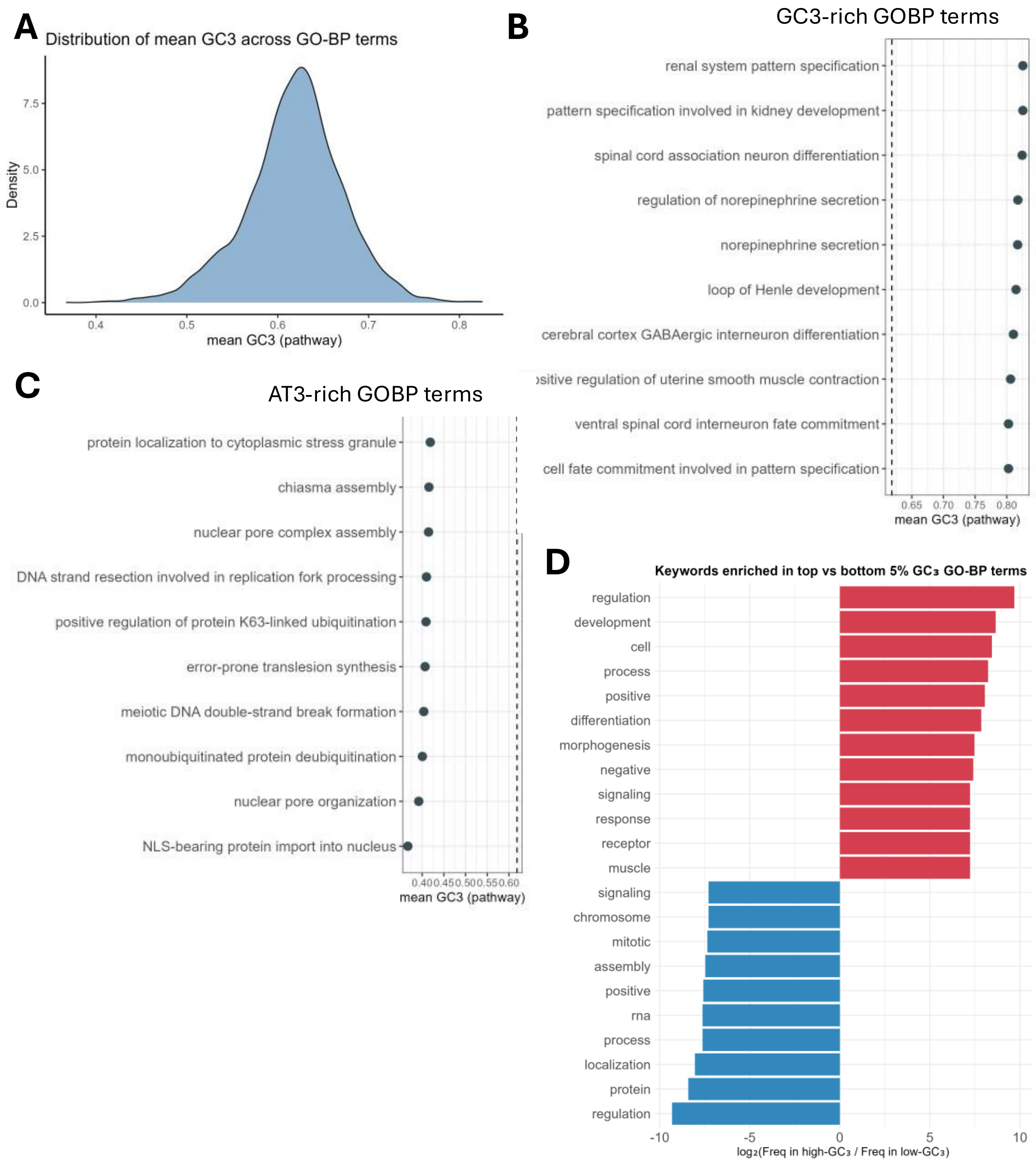
A
Distribution of mean GC3 across GO-BP terms
Density
mean GC3 (pathway)
B
GC3-rich GOBP terms
renal system pattern specification
pattern specification involved in kidney development
spinal cord association neuron differentiation
regulation of norepinephrine secretion
norepinephrine secretion
loop of Henle development
cerebral cortex GABAergic interneuron differentiation
ventral spinal cord interneuron fate commitment
cell fate commitment involved in pattern specification
mean GC3 (pathway)
C
AT3-rich GOBP terms
protein localization to cytoplasmic stress granule
chiasma assembly
nuclear pore complex assembly
DNA strand resection involved in replication fork processing
positive regulation of protein K63-linked ubiquitination
error-prone translesion synthesis
meiotic DNA double-strand break formation
monoubiquitinated protein deubiquitination
nuclear pore organization
NLS-bearing protein import into nucleus
mean GC3 (pathway)
D
Keywords enriched in top vs bottom 5% GC3 GO-BP terms
regulation
development
cell
process
positive
differentiation
morphogenesis
negative
signaling
response
receptor
muscle
signaling
chromosome
mitotic
assembly
positive
rna
process
localization
protein
regulation
log2(Freq in high-GC3 / Freq in low-GC3)

**Supplementary figure 5**: **Isoacceptors frequencies analysis of GOBP Terms. A:** Heatmap of isoacceptors frequencies T-stats of all tested GOBP terms (T-stat represents the results of T-test of the isoacceptors frequencies of genes present in a GOBP term versus the background, i.e., all genes). **B:** Heatmap showing isoacceptors frequencies T-stat Top G/C-ending biased GOBP pathways. **C:** Heatmap showing isoacceptors frequencies T-stat Top A/T-ending biased GOBP pathways.

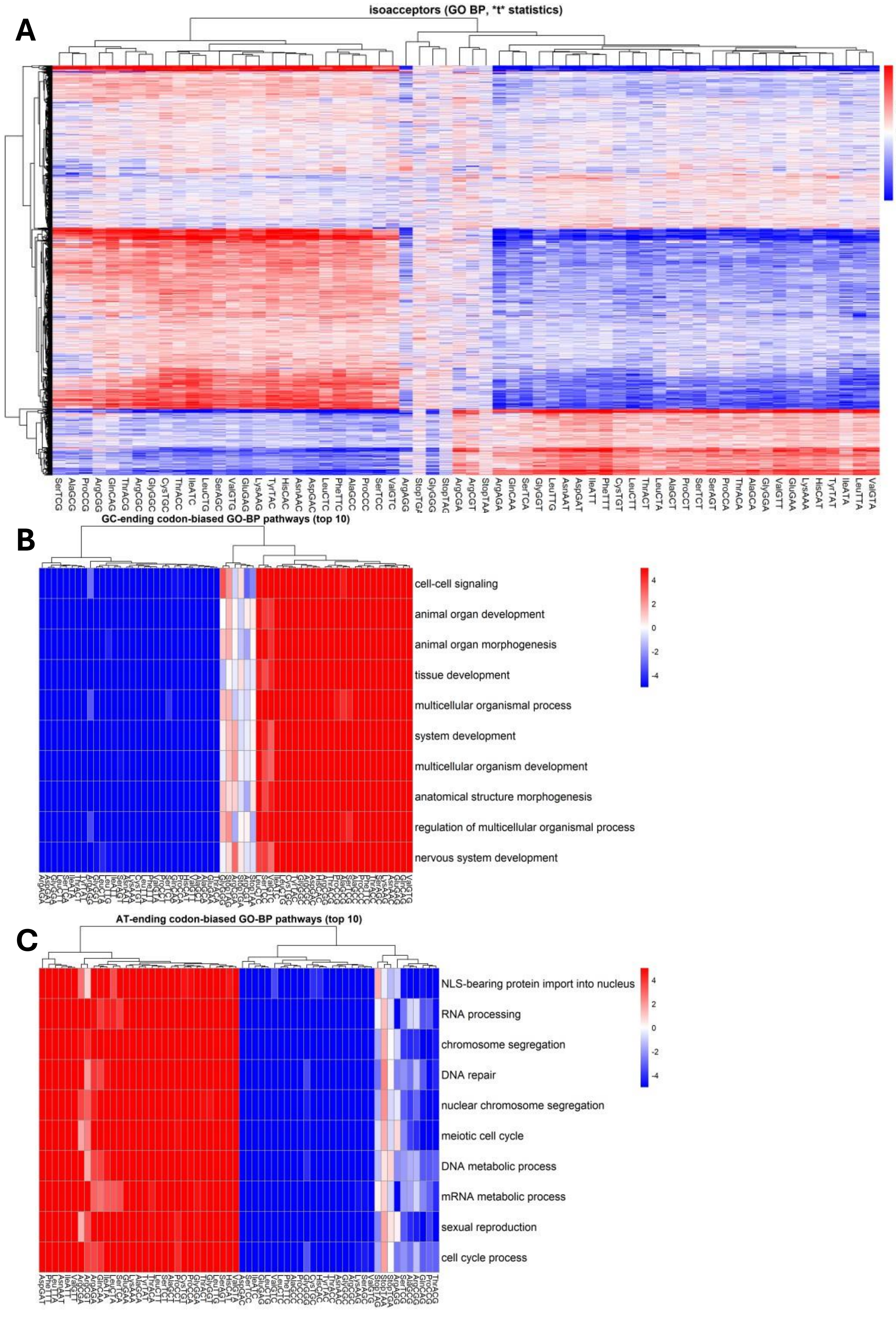
A
isoacceptors (GO BP, *t* statistics)
4
2
0
-2
-4
SerTCG AlaGCG ProCCG ArgCGG GlnCAG ThrACG ArgCGC GlyGGC CysTGC ThrACC IleATC LeuCTG SerAGC ValGTG GluGAG LysAAG TyrTAC HisCAC AsnAAC AspGAC LeuCTC PheTTC AlaGCC ProCCC SerTCC ValGTC ArgAGG StopTGA GlyGGG StopTAG ArgCGA ArgCGT StopTAA ArgAGA GlnCAA SerTCA GlyGGT LeuTTG AsnAAT AspGAT IleATT PheTTT CysTGT LeuCTT ThrACT LeuCTA AlaGCT ProCCT SerTCT SerAGT ProCCA ThrACA AlaGCA GlyGGA ValGTT GluGAA LysAAA HisCAT TyrTAT IleATA LeuTTA ValGTA
B
GC-ending codon-biased GO-BP pathways (top 10)
cell-cell signaling
animal organ development
animal organ morphogenesis
tissue development
multicellular organismal process
system development
multicellular organism development
anatomical structure morphogenesis
regulation of multicellular organismal process
nervous system development
4
2
0
-2
-4
C
AT-ending codon-biased GO-BP pathways (top 10)
NLS-bearing protein import into nucleus
RNA processing
chromosome segregation
DNA repair
nuclear chromosome segregation
meiotic cell cycle
DNA metabolic process
mRNA metabolic process
sexual reproduction
cell cycle process
4
2
0
-2
-4

**Supplementary figure 6: Analysis of functional enrichment of leucine synonymous codons.** **A:** density plot of Leu codons isoacceptors frequencies across species. **B:** ORA GOBP of the top genes biased towards each of the Leu synonymous codons.

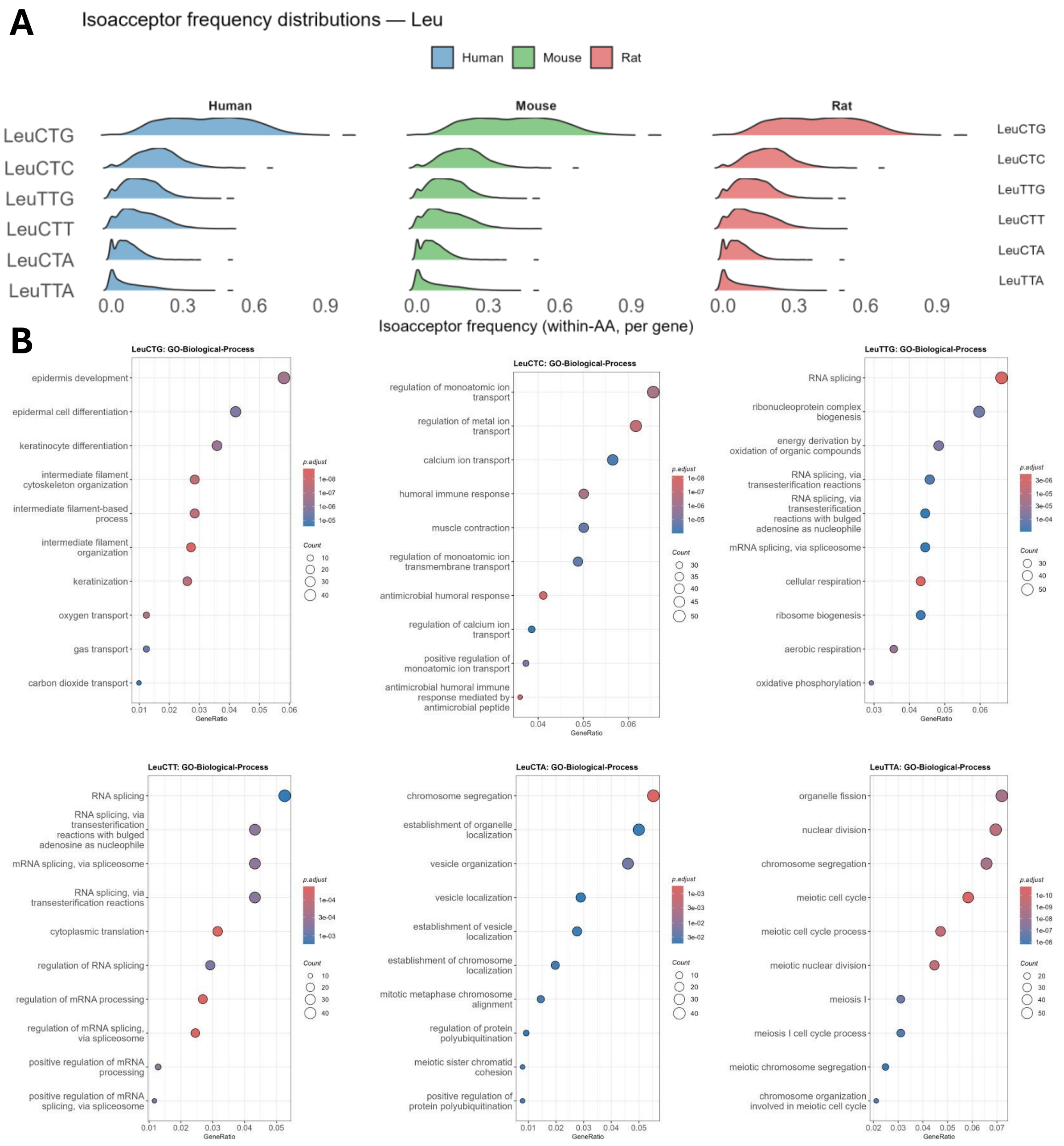
A
Isoacceptor frequency distributions — Leu
Human
Mouse
Rat
LeuCTG
LeuCTC
LeuTTG
LeuCTT
LeuCTA
LeuTTA
0.0
0.3
0.6
0.9
Isoacceptor frequency (within-AA, per gene)
B
LeuCTG: GO-Biological-Process
epidermis development
epidermal cell differentiation
keratinocyte differentiation
intermediate filament cytoskeleton organization
intermediate filament-based process
intermediate filament organization
keratinization
oxygen transport
gas transport
carbon dioxide transport
GeneRatio
p.adjust
Count
LeuCTC: GO-Biological-Process
regulation of monoatomic ion transport
regulation of metal ion transport
calcium ion transport
humoral immune response
muscle contraction
regulation of monoatomic ion transmembrane transport
antimicrobial humoral response
regulation of calcium ion transport
positive regulation of monoatomic ion transport
antimicrobial humoral immune response mediated by antimicrobial peptide
LeuTTG: GO-Biological-Process
RNA splicing
ribonucleoprotein complex biogenesis
energy derivation by oxidation of organic compounds
RNA splicing, via transesterification reactions
RNA splicing, via transesterification reactions with bulged adenosine as nucleophile
mRNA splicing, via spliceosome
cellular respiration
ribosome biogenesis
aerobic respiration
oxidative phosphorylation
LeuCTT: GO-Biological-Process
RNA splicing
RNA splicing, via transesterification reactions with bulged adenosine as nucleophile
mRNA splicing, via spliceosome
RNA splicing, via transesterification reactions
cytoplasmic translation
regulation of RNA splicing
regulation of mRNA processing
regulation of mRNA splicing, via spliceosome
positive regulation of mRNA processing
positive regulation of mRNA splicing, via spliceosome
LeuCTA: GO-Biological-Process
chromosome segregation
establishment of organelle localization
vesicle organization
vesicle localization
establishment of vesicle localization
establishment of chromosome localization
mitotic metaphase chromosome alignment
regulation of protein polyubiquitination
meiotic sister chromatid cohesion
positive regulation of protein polyubiquitination
LeuTTA: GO-Biological-Process
organelle fission
nuclear division
chromosome segregation
meiotic cell cycle
meiotic cell cycle process
meiotic nuclear division
meiosis I
meiosis I cell cycle process
meiotic chromosome segregation
chromosome organization involved in meiotic cell cycle

**Supplementary figure 7: Analysis of ANN-index and its links to cancer oncogenesis. A:** ANN-index density plot across human genes. **B:** GOBP ORA of top 10% genes by ANN-index scores. **C:** GOBP ORA of Bottom 10% genes by ANN-index scores. **D:** Spearman's rank correlation analysis between gene ANN-index and GC3 scores.

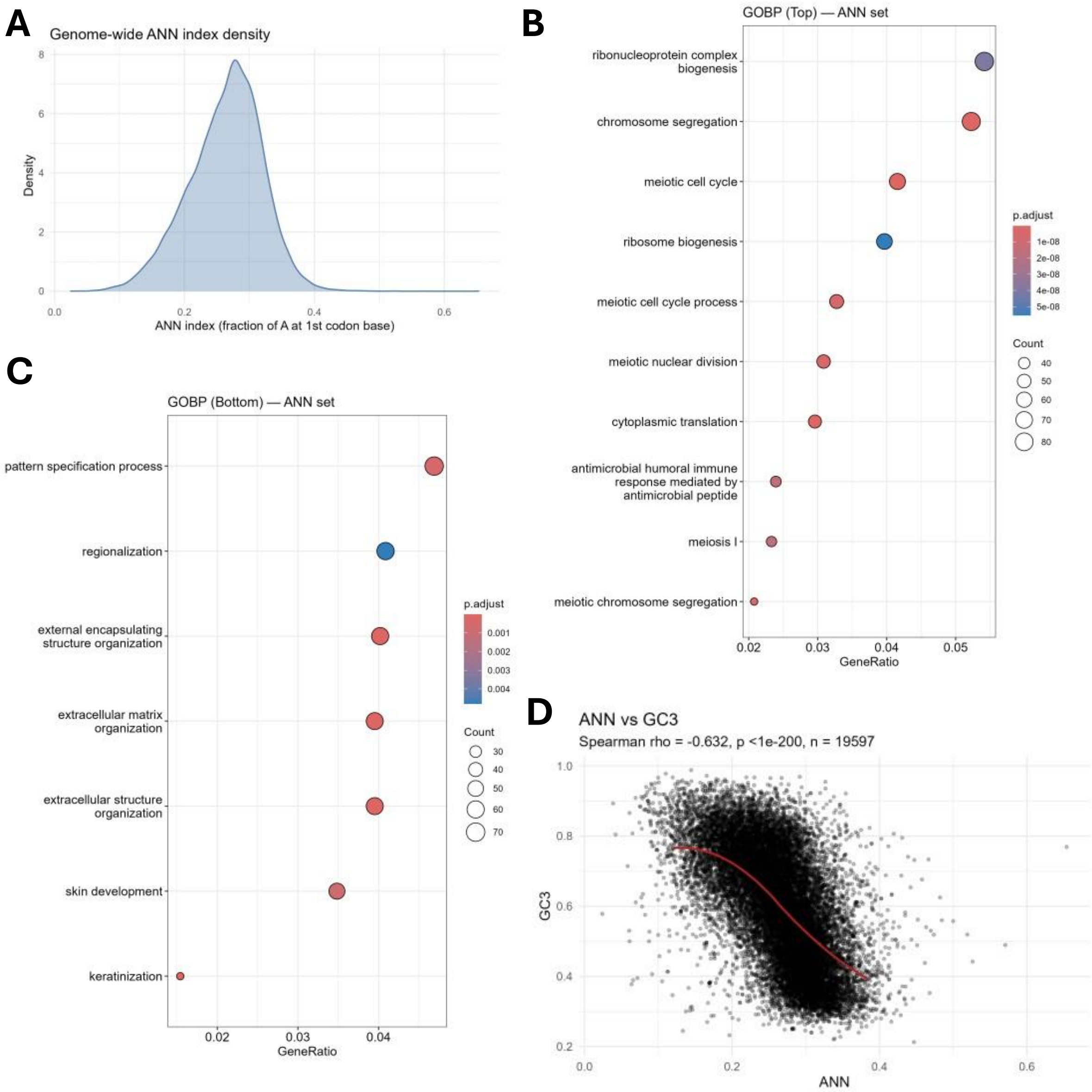
A
Genome-wide ANN index density
Density
ANN index (fraction of A at 1st codon base)
B
GOBP (Top) — ANN set
ribonucleoprotein complex biogenesis
chromosome segregation
meiotic cell cycle
ribosome biogenesis
meiotic cell cycle process
meiotic nuclear division
cytoplasmic translation
antimicrobial humoral immune response mediated by antimicrobial peptide
meiosis I
meiotic chromosome segregation
GeneRatio
p.adjust
Count
C
GOBP (Bottom) — ANN set
pattern specification process
regionalization
external encapsulating structure organization
extracellular matrix organization
extracellular structure organization
skin development
keratinization
GeneRatio
p.adjust
Count
D
ANN vs GC3
Spearman rho = -0.632, p <1e-200, n = 19597
GC3
ANN

**Supplementary figure 8: Analysis of m$^7$G-index and its links to cancer oncogenesis. A:** m$^7$G-index density plot across human genes. **B:** GOBP ORA of top 10% genes by m$^7$G-index scores. **C:** GOBP ORA of Bottom 10% genes by m$^7$G-index scores. **D:** Spearman's rank correlation analysis between gene m$^7$G-index and GC3 scores. **E:** Spearman's rank correlation analysis between gene m$^7$G-index and ANN-index scores.

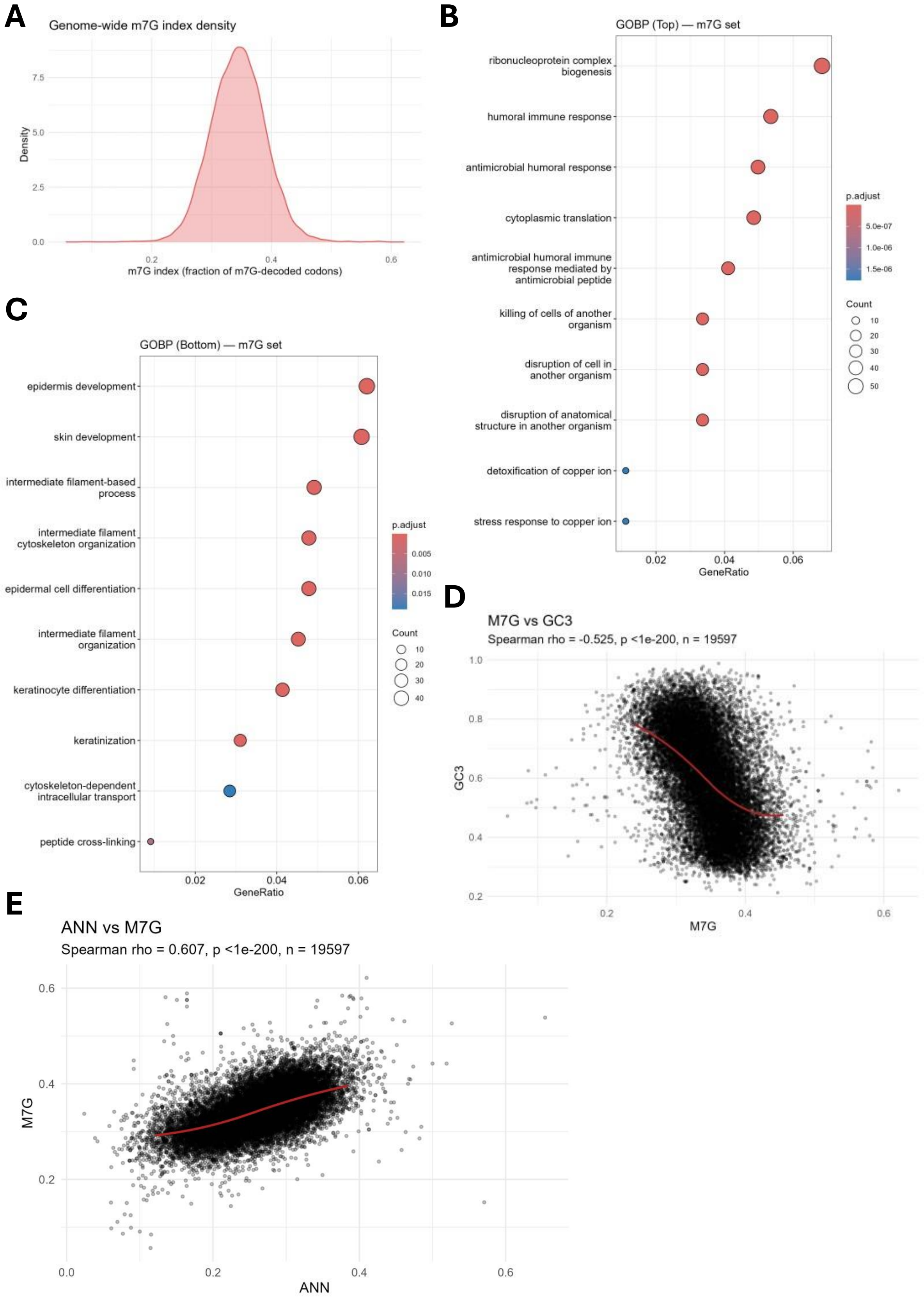
A
Genome-wide m7G index density
Density
7.5
5.0
2.5
0.0
0.2
0.4
0.6
m7G index (fraction of m7G-decoded codons)
B
GOBP (Top) — m7G set
ribonucleoprotein complex biogenesis
humoral immune response
antimicrobial humoral response
cytoplasmic translation
antimicrobial humoral immune response mediated by antimicrobial peptide
killing of cells of another organism
disruption of cell in another organism
disruption of anatomical structure in another organism
detoxification of copper ion
stress response to copper ion
0.02
0.04
0.06
GeneRatio
p.adjust
5.0e-07
1.0e-06
1.5e-06
Count
10
20
30
40
50
C
GOBP (Bottom) — m7G set
epidermis development
skin development
intermediate filament-based process
intermediate filament cytoskeleton organization
epidermal cell differentiation
intermediate filament organization
keratinocyte differentiation
keratinization
cytoskeleton-dependent intracellular transport
peptide cross-linking
0.02
0.04
0.06
GeneRatio
p.adjust
0.005
0.010
0.015
Count
10
20
30
40
D
M7G vs GC3
Spearman rho = -0.525, p <1e-200, n = 19597
GC3
1.0
0.8
0.6
0.4
0.2
0.2
0.4
0.6
M7G
E
ANN vs M7G
Spearman rho = 0.607, p <1e-200, n = 19597
M7G
0.6
0.4
0.2
0.0
0.2
0.4
0.6
ANN

**Supplementary figure 9: GOBP ORA analysis of human tissues expressed proteins. A:** Heatmap GOBP ORA enrichment across different human tissues. **B:** GOBP ORA analysis of the top 10% expressed proteins in Testis. **C:** GOBP ORA analysis of the top 10% expressed proteins in Cerebellum.

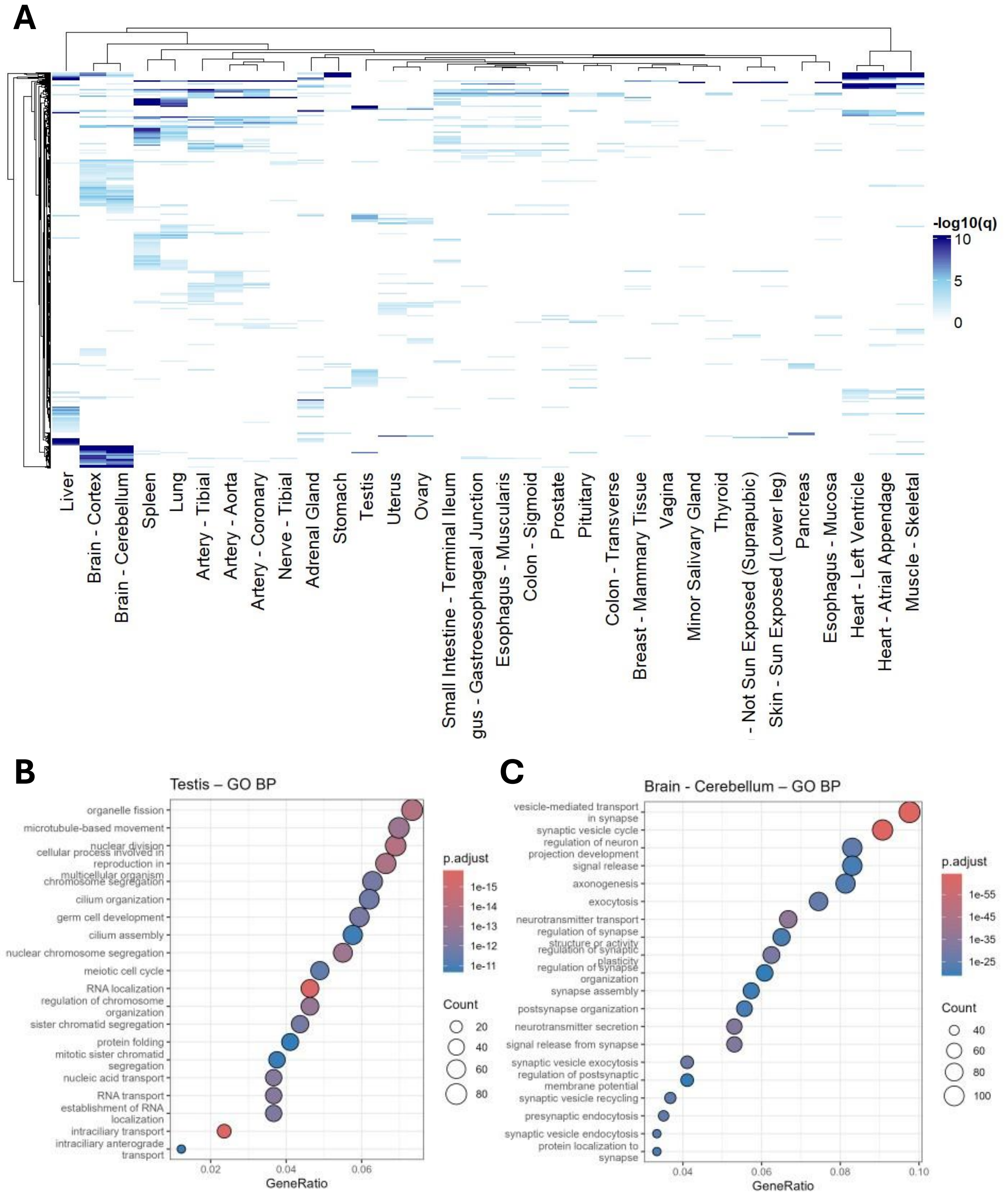

A
-log10(q)
10
5
0
Liver
Brain - Cortex
Brain - Cerebellum
Spleen
Lung
Artery - Tibial
Artery - Aorta
Artery - Coronary
Nerve - Tibial
Adrenal Gland
Stomach
Testis
Uterus
Ovary
Small Intestine - Terminal Ileum
gus - Gastroesophageal Junction
Esophagus - Muscularis
Colon - Sigmoid
Prostate
Pituitary
Colon - Transverse
Breast - Mammary Tissue
Vagina
Minor Salivary Gland
Thyroid
- Not Sun Exposed (Suprapubic)
Skin - Sun Exposed (Lower leg)
Pancreas
Esophagus - Mucosa
Heart - Left Ventricle
Heart - Atrial Appendage
Muscle - Skeletal
B
Testis – GO BP
organelle fission
microtubule-based movement
nuclear division
cellular process involved in reproduction in multicellular organism
chromosome segregation
cilium organization
germ cell development
cilium assembly
nuclear chromosome segregation
meiotic cell cycle
RNA localization
regulation of chromosome organization
sister chromatid segregation
protein folding
mitotic sister chromatid segregation
nucleic acid transport
RNA transport
establishment of RNA localization
intraciliary transport
intraciliary anterograde transport
0.02
0.04
0.06
GeneRatio
p.adjust
1e-15
1e-14
1e-13
1e-12
1e-11
Count
20
40
60
80
C
Brain - Cerebellum – GO BP
vesicle-mediated transport in synapse
synaptic vesicle cycle
regulation of neuron projection development
signal release
axonogenesis
exocytosis
neurotransmitter transport
regulation of synapse structure or activity
regulation of synaptic plasticity
regulation of synapse organization
synapse assembly
postsynapse organization
neurotransmitter secretion
signal release from synapse
synaptic vesicle exocytosis
regulation of postsynaptic membrane potential
synaptic vesicle recycling
presynaptic endocytosis
synaptic vesicle endocytosis
protein localization to synapse
0.04
0.06
0.08
0.10
GeneRatio
p.adjust
1e-55
1e-45
1e-35
1e-25
Count
40
60
80
100

**Supplementary figure 10:** **A:** Density plot of mean GC3 scores of the top 20 GOBP terms enriched in each tissue. **B:** Density plot showing the mean GC3 scores of the top 20 GOBP terms in the top and bottom 3 tissues by protein GC3 scores.

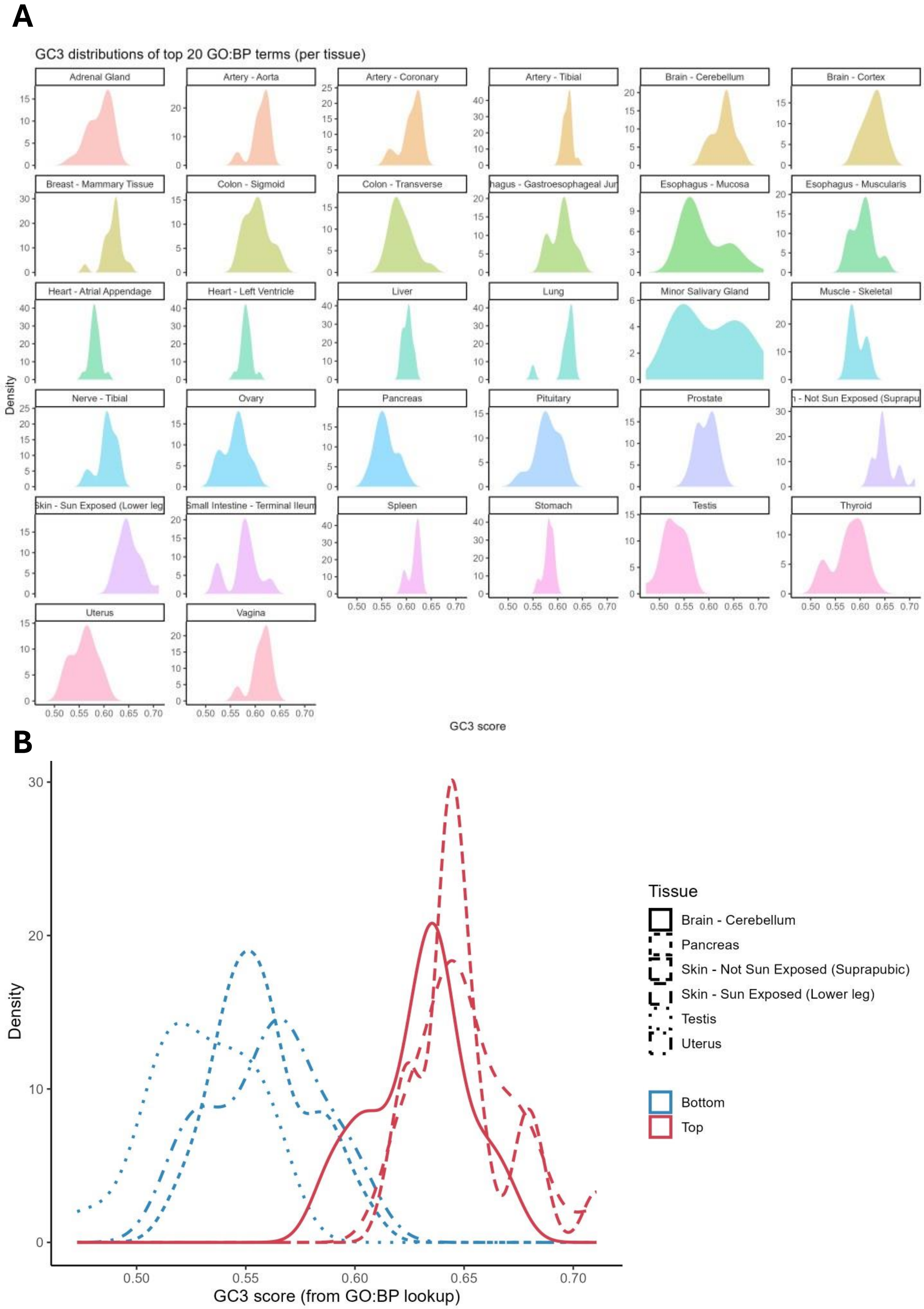
A
GC3 distributions of top 20 GO:BP terms (per tissue)
Adrenal Gland
Artery - Aorta
Artery - Coronary
Artery - Tibial
Brain - Cerebellum
Brain - Cortex
Breast - Mammary Tissue
Colon - Sigmoid
Colon - Transverse
hagus - Gastroesophageal Jun
Esophagus - Mucosa
Esophagus - Muscularis
Heart - Atrial Appendage
Heart - Left Ventricle
Liver
Lung
Minor Salivary Gland
Muscle - Skeletal
Nerve - Tibial
Ovary
Pancreas
Pituitary
Prostate
- Not Sun Exposed (Suprapu
Skin - Sun Exposed (Lower leg
Small Intestine - Terminal Ileum
Spleen
Stomach
Testis
Thyroid
Uterus
Vagina
Density
GC3 score
B
Density
GC3 score (from GO:BP lookup)
Tissue
Brain - Cerebellum
Pancreas
Skin - Not Sun Exposed (Suprapubic)
Skin - Sun Exposed (Lower leg)
Testis
Uterus
Bottom
Top

**Supplementary figure 11:** Cell models passports database quality control and analytics. **A:** Bar plot showing the number of models per tissue of origin. **B:** Bar plot showing the number of models per tissue status (tumor, normal, metastasis, etc.). **C:** Histogram of the number of detected RNAs per model across all models (1 ⩾ TPM). **D:** Histogram of the number of detected proteins per model across all models.

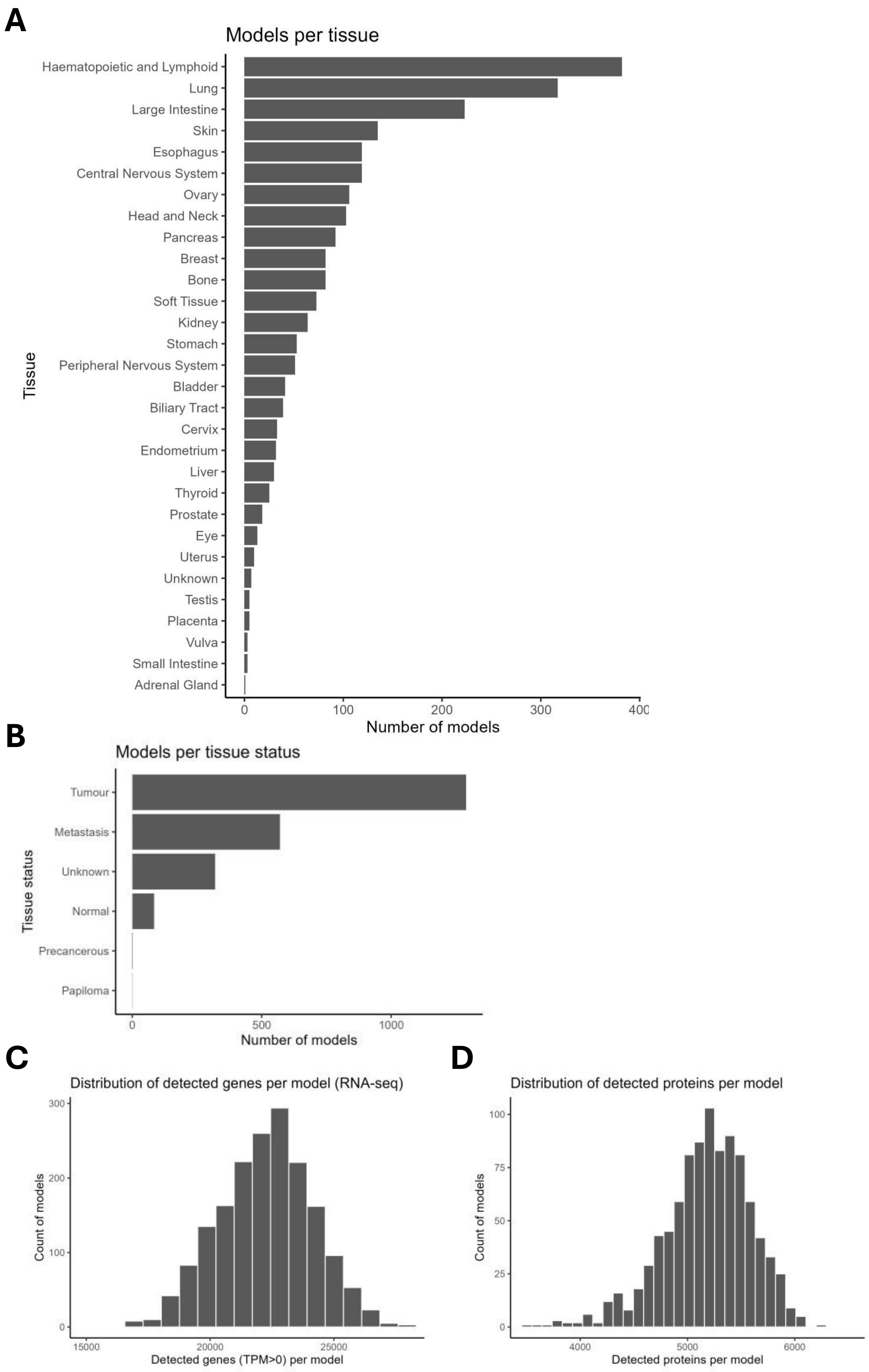
A
Models per tissue
Haematopoietic and Lymphoid
Lung
Large Intestine
Skin
Esophagus
Central Nervous System
Ovary
Head and Neck
Pancreas
Breast
Bone
Soft Tissue
Kidney
Stomach
Peripheral Nervous System
Bladder
Biliary Tract
Cervix
Endometrium
Liver
Thyroid
Prostate
Eye
Uterus
Unknown
Testis
Placenta
Vulva
Small Intestine
Adrenal Gland
Tissue
0 100 200 300 400
Number of models
B
Models per tissue status
Tumour
Metastasis
Unknown
Normal
Precancerous
Papiloma
Tissue status
0 500 1000
Number of models
C
Distribution of detected genes per model (RNA-seq)
Count of models
0 100 200 300
15000 20000 25000
Detected genes (TPM>0) per model
D
Distribution of detected proteins per model
Count of models
0 25 50 75 100
4000 5000 6000
Detected proteins per model

**Supplementary figure 12: A:** Alluvial clustering analysis of the K-mean clusters in Figure 8C showing the membership and features of different models in each cluster. **B:** Heatmap of isoacceptors frequencies T-stat of cell lines from central nervous system tumors. **C:** Heatmap of isoacceptors frequencies T-stat of cell lines from hematopoietic and lymphoid tumors.

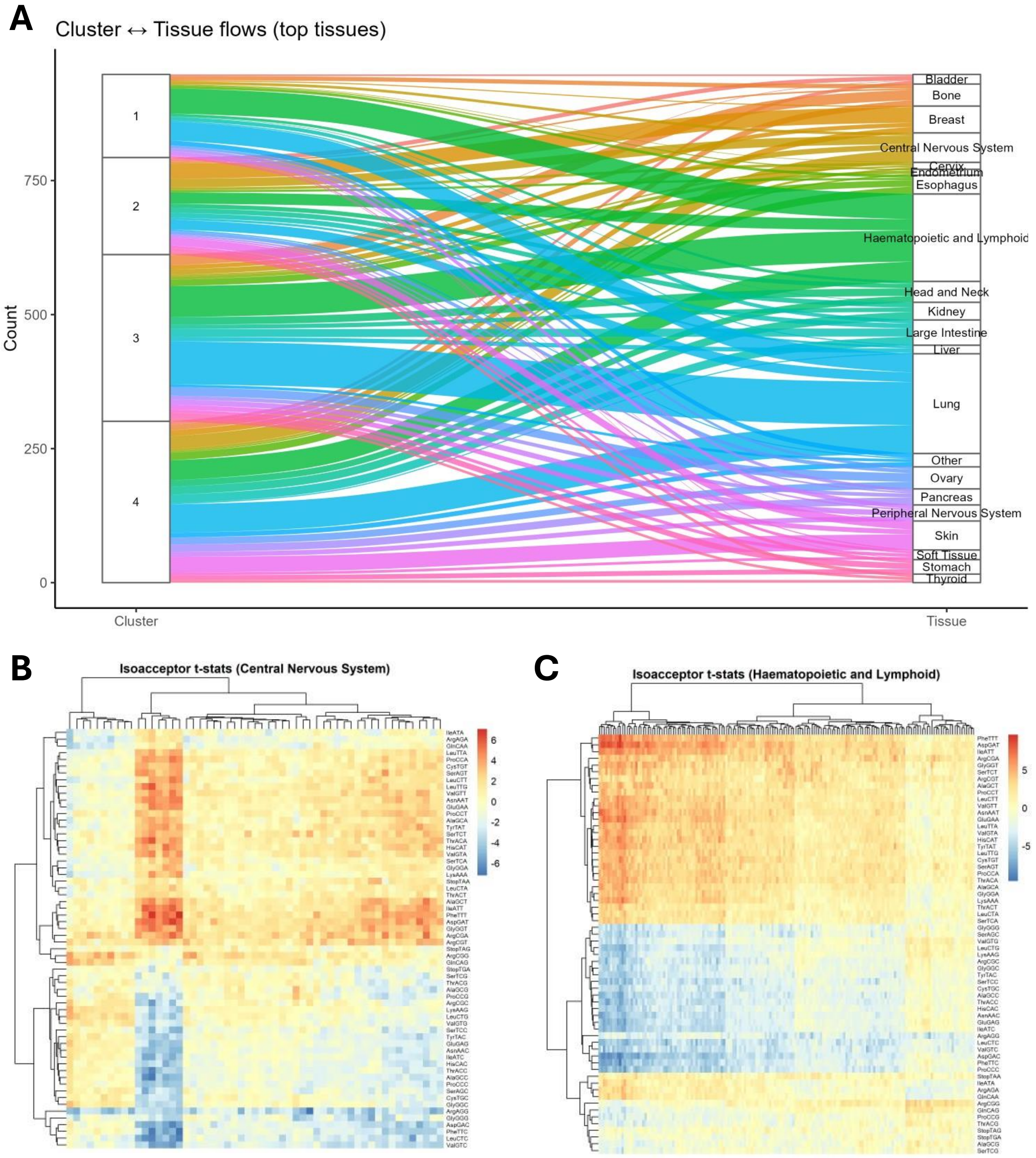
A
Cluster ↔ Tissue flows (top tissues)
Count
0
250
500
750
1
2
3
4
Bladder
Bone
Breast
Central Nervous System
Cervix
Endometrium
Esophagus
Haematopoietic and Lymphoid
Head and Neck
Kidney
Large Intestine
Liver
Lung
Other
Ovary
Pancreas
Peripheral Nervous System
Skin
Soft Tissue
Stomach
Thyroid
Cluster
Tissue
B
Isoacceptor t-stats (Central Nervous System)
6
4
2
0
-2
-4
-6
C
Isoacceptor t-stats (Haematopoietic and Lymphoid)
5
0
-5

**Supplementary figure 13:** Analysis of model features in K-mean clusters. **A:** Representation of different tissues of origin in each K-mean cluster. **B:** Cancer type representation in each cluster. **C:** Tissue status representation in each cluster. **D:** Cell models growth properties representation in each cluster.

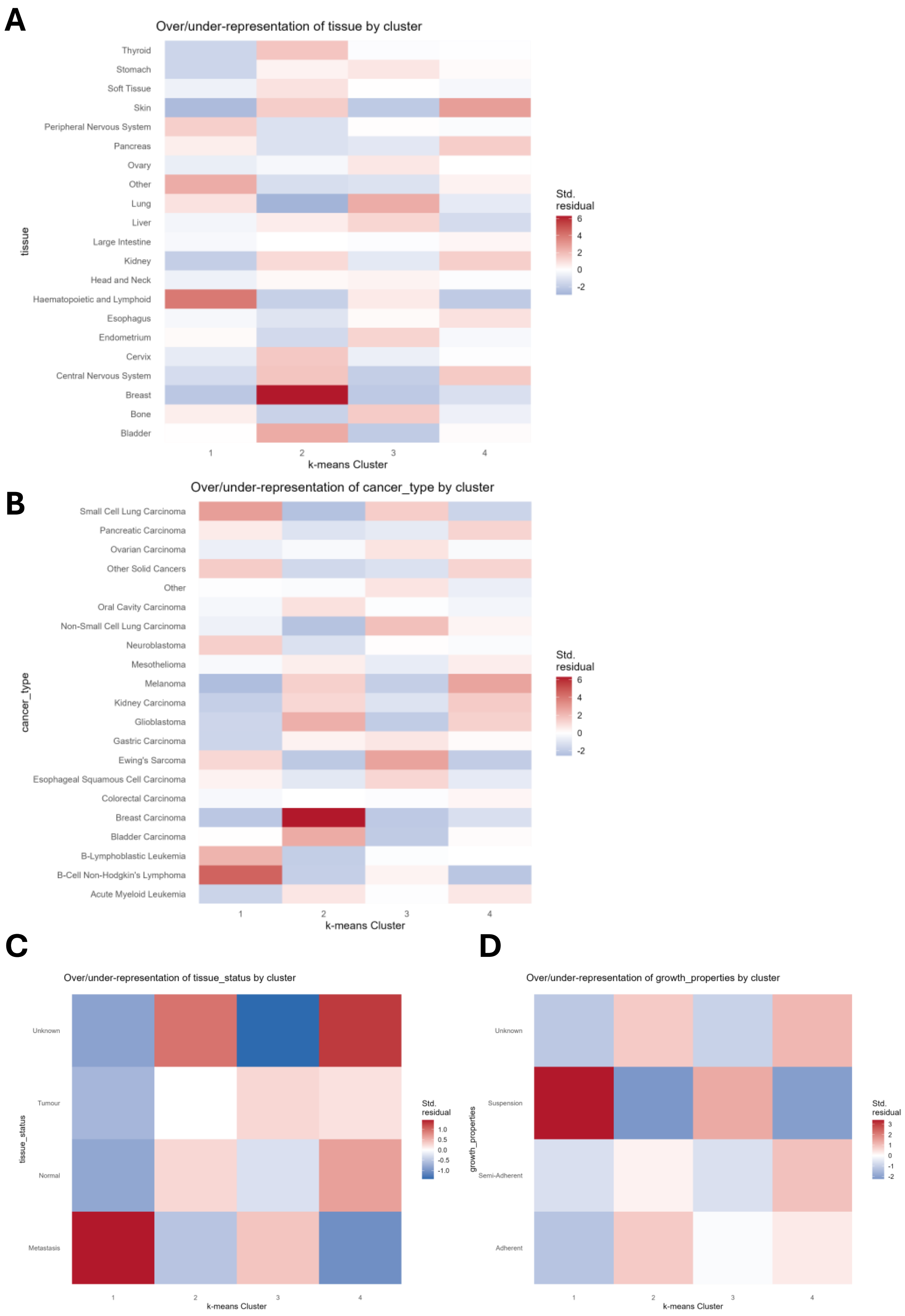
A
Over/under-representation of tissue by cluster
Thyroid
Stomach
Soft Tissue
Skin
Peripheral Nervous System
Pancreas
Ovary
Other
Lung
Liver
Large Intestine
Kidney
Head and Neck
Haematopoietic and Lymphoid
Esophagus
Endometrium
Cervix
Central Nervous System
Breast
Bone
Bladder
tissue
1
2
3
4
k-means Cluster
Std. residual
6
4
2
0
-2
B
Over/under-representation of cancer_type by cluster
Small Cell Lung Carcinoma
Pancreatic Carcinoma
Ovarian Carcinoma
Other Solid Cancers
Other
Oral Cavity Carcinoma
Non-Small Cell Lung Carcinoma
Neuroblastoma
Mesothelioma
Melanoma
Kidney Carcinoma
Glioblastoma
Gastric Carcinoma
Ewing's Sarcoma
Esophageal Squamous Cell Carcinoma
Colorectal Carcinoma
Breast Carcinoma
Bladder Carcinoma
B-Lymphoblastic Leukemia
B-Cell Non-Hodgkin's Lymphoma
Acute Myeloid Leukemia
cancer_type
1
2
3
4
k-means Cluster
Std. residual
6
4
2
0
-2
C
Over/under-representation of tissue_status by cluster
Unknown
Tumour
Normal
Metastasis
tissue_status
1
2
3
4
k-means Cluster
Std. residual
1.0
0.5
0.0
-0.5
-1.0
D
Over/under-representation of growth_properties by cluster
Unknown
Suspension
Semi-Adherent
Adherent
growth_properties
1
2
3
4
k-means Cluster
Std. residual
3
2
1
0
-1
-2

**Supplementary figure 14: Human cancers isoacceptors codon frequencies are homogeneous across patients.** Analysis of the isoacceptors frequencies of the top 10% proteins expressed in human cancer samples from Knol et al study (Cancer cell, 2025) reveals homogeneous codon usage across samples belonging to the same tumor type.

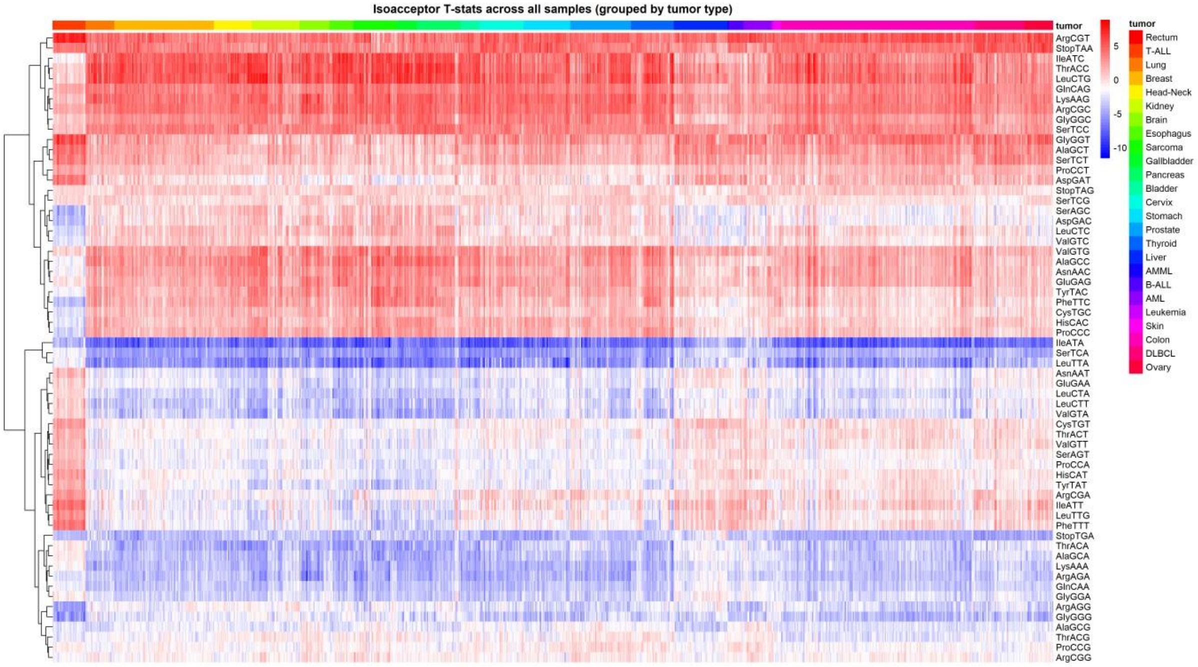

Isoacceptor T-stats across all samples (grouped by tumor type)
tumor
ArgCGT
StopTAA
IleATC
ThrACC
LeuCTG
GlnCAG
LysAAG
ArgCGC
GlyGGC
SerTCC
GlyGGT
AlaGCT
SerTCT
ProCCT
AspGAT
StopTAG
SerTCG
SerAGC
AspGAC
LeuCTC
ValGTC
ValGTG
AlaGCC
AsnAAC
GluGAG
TyrTAC
PheTTC
CysTGC
HisCAC
ProCCC
IleATA
SerTCA
LeuTTA
AsnAAT
GluGAA
LeuCTA
LeuCTT
ValGTA
CysTGT
ThrACT
ValGTT
SerAGT
ProCCA
HisCAT
TyrTAT
ArgCGA
IleATT
LeuTTG
PheTTT
StopTGA
ThrACA
AlaGCA
LysAAA
ArgAGA
GlnCAA
GlyGGA
ArgAGG
GlyGGG
AlaGCG
ThrACG
ProCCG
ArgCGG
5
0
-5
-10
tumor
Rectum
T-ALL
Lung
Breast
Head-Neck
Kidney
Brain
Esophagus
Sarcoma
Gallbladder
Pancreas
Bladder
Cervix
Stomach
Prostate
Thyroid
Liver
AMML
B-ALL
AML
Leukemia
Skin
Colon
DLBCL
Ovary

**Supplementary figure 15: Analysis of human cancer proteomes compared to normal tissues**. **A:** Global GC3 scores of upregulated and downregulated proteins ($|log2FC| > 0.58$ and $FDR < 0.05$) across all cancers compared to their respective normal tissues. **B:** Global GC3 scores of upregulated and downregulated proteins ($|log2FC| > 0.58$ and $FDR < 0.05$) in acute myeloid leukemia. **C:** Heatmap of different cancers isoacceptors frequencies. Upregulated and downregulated proteins were analyzed separately and compared against the genome to generate T-statistics which were used for visualization. **D:** Cancer abbreviations and full names.

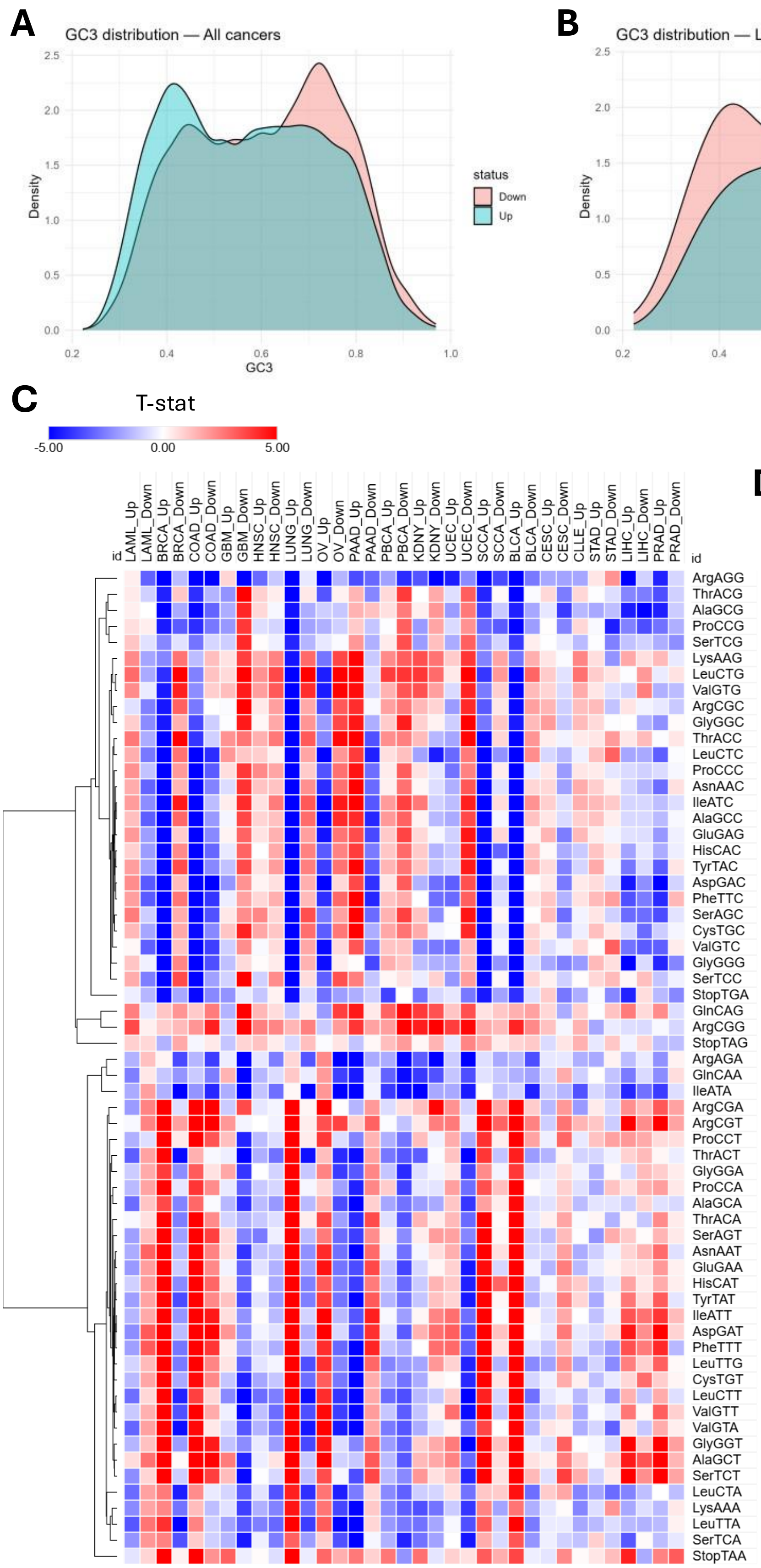


**D**

| Abbreviation | Full name |
|---|---|
| SCCA | Anal Canal Cancer |
| LAML | Acute Myeloid Leukemia |
| CLLE | Chronic Lymphocytic Leukemia |
| BRCA | Breast Cancer |
| BLCA | Bladder Cancer |
| KDNY | Kidney Cancer |
| COAD | Colon Cancer |
| STAD | Gastric Cancer |
| HNSC | Head and Neck Squamous Cell Carcinoma |
| LIHC | Liver Cancer |
| LUNG | Lung Cancer |
| ORCA | Oral Squamous Cell Carcinoma |
| OV | Ovary Cancer |
| PAAD | Pancreatic Cancer |
| GBM | Glioblastoma |
| PBCA | PediatricAYA Brain Tumors |
| UCEC | Endometrial Carcinoma |
| PRAD | Prostate Cancer |
| SKCM | Skin Cancer |
| THCA | Thyroid Cancer |
| CESC | Cervical Cancer |
